# High resolution mouse subventricular zone stem cell niche transcriptome reveals features of lineage, anatomy, and aging

**DOI:** 10.1101/2020.07.27.223602

**Authors:** Xuanhua P. Xie, Dan R. Laks, Daochun Sun, Asaf Poran, Ashley M. Laughney, Zilai Wang, Jessica Sam, German Belenguer, Isabel Fariñas, Olivier Elemento, Xiuping Zhou, Luis F. Parada

**Affiliations:** Cancer Biology and Genetics Program; Brain Tumor Center, Memorial Sloan Kettering Cancer Center, 1275 York Ave., New York, NY 10065, USA; Institute for Computational Biomedicine, Department of Physiology and Biophysics; Weill Cornell Medicine, New York, NY 10065, USA; Biochemistry, Cell & Molecular Biology Graduate Program, Weill Cornell Medicine, New York, NY 10065, USA; Centro de Investigación Biomédica en Red sobre Enfermedades Neurodegenerativas (CIBERNED), Madrid 28031, Spain; Departamento de Biología Celular, Biología Funcional y Antropología Física, Universidad de Valencia, Valencia 46010, Spain; Institute of Nervous System Diseases, Xuzhou Medical University, Jiangsu 221002, PR China; Neon Therapeutics, Cambridge, Massachusetts, MA, USA

**Keywords:** subventricular zone (SVZ), neural stem cell (NSC), single cell RNA sequencing, transcriptome, aging

## Abstract

Adult neural stem cells (NSC) serve as a reservoir for brain plasticity and origin for certain gliomas. Lineage tracing and genomic approaches have portrayed complex underlying heterogeneity within the major anatomical location for NSC, the subventricular zone (SVZ). To gain a comprehensive profile of NSC heterogeneity, we utilized a well validated stem/progenitor specific reporter transgene in concert with single cell RNA sequencing to achieve unbiased analysis of SVZ cells from infancy to advanced age. The magnitude and high specificity of the resulting transcriptional data sets allow precise identification of the varied cell types embedded in the SVZ including specialized parenchymal cells (neurons, glia, microglia), and non-central nervous system cells (endothelial, immune). Initial mining of the data delineates four quiescent NSC and three progenitor cell subpopulations formed in a linear progression. Further evidence indicates that distinct stem and progenitor populations reside in different regions of the SVZ. As stem/progenitor populations progress from neonatal to advanced age, they acquire a deficiency in transition from quiescence to proliferation. Further data mining identifies stage specific biological processes, transcription factor networks, and cell surface markers for investigation of cellular identities, lineage relationships, and key regulatory pathways in adult NSC maintenance and neurogenesis.

**Significance Statement:** Adult neural stem cells (NSC) are closely related to multiple neurological disorders and brain tumors. Comprehensive investigation of their composition, lineage, and aging will provide new insights that may lead to enhanced patient treatment. This study applies a novel transgene to label and manipulate neural stem/progenitor cells, and monitor their evolution during aging. Together with high-throughput single cell RNA sequencing, we are able to analyze the subventricular zone (SVZ) cells from infancy to advanced age with unprecedented granularity. Diverse new cell states are identified in the stem cell niche, and an aging related NSC deficiency in transition from quiescence to proliferation is identified. The related biological features provide rich resources to inspect adult NSC maintenance and neurogenesis.

## Introduction

Most mammalian organs and tissues harbor stem cells that serve as a source of continual cell replenishment. Adult rat neurogenesis was first documented in the 1960’s and since then the rodent brain has served as a useful model for study of neural stem cells (NSC) (1). In the mouse brain, two distinct pools of NSC, found in the subventricular zone (SVZ) of the lateral ventricle walls and in the dentate gyrus of the hippocampus, produce new neurons that functionally integrate into pre-existing circuits during lifetime (2–6). In the SVZ, both type B1 cells and ependymal cells had been proposed as NSC (7–10). B1 cells are elongated stem cells that belong to the radial glia-astrocytic lineage and express markers characteristically found in astrocytes, such as glial fibrillary acidic protein (GFAP) or glutamate-aspartate transporter (GLAST-Slc1a3). They exhibit a thin cytoplasmic process that intercalates between ependymal cells to adorn the ventricle cavities and terminates in a sensing primary cilium and another long process that interacts with the basal lamina of irrigating vasculature. B1 cells give rise to migratory and dividing transient amplifying progenitor (TAP) cells that differentiate into interneurons destined to the olfactory bulb (OB) or, less frequently, into oligodendrocytes in the corpus callosum (11, 12).

Apart from their role in OB interneuron replacement and to a lesser degree, to other neurogenesis and gliogenesis, it has been reported that SVZ NSC/progenitor cells can mount responses to brain cell death caused by trauma (13, 14). In addition, increasing evidence points to NSC as a relevant source for malignant transformation into glioma. Direct oncogenic transformation of mouse NSC or progenitors has been demonstrated to mediate GBM and medulloblastoma (15, 16). Mouse modeling studies have portrayed a direct lineage relationship between NSC/OLC (Oligodendrocyte lineage cell) stem/progenitor cells and GBM (17, 18). Thus, a complete understanding of the properties of SVZ NSC, their transition into TAPs, and finally into differentiated cells, will hold relevance to brain physiology and repair, and to the etiology of malignant brain tumors.

Advances in genomics, deep sequencing, and microfluidics have allowed increased resolution of the SVZ stem cell niche cell dynamics (6, 10, 19–24). NSC surface markers, such as Prom1-CD133 and Slc1a3-Glast, and transgenes encoding Cre-recombinase or fluorescent reporters regulated by Gfap promoters, have been used to partially enrich for prospective NSC from the SVZ for bulk RNA sequencing (RNAseq). The data yielded information consistent with the existence of a quiescent NSC population that transitions to an activated state (19–22). Single cell transcriptomics of SVZ derived cells have further enhanced resolution. Recent studies relying on cell sorting with NSC or TAP markers have yielded limited numbers of cells for single cell sequencing ranging from tens to hundreds of cells (10, 23, 24).

To more efficiently label and study NSC, we constructed a novel transgene with a GFP reporter that highlights the vast majority of stem and progenitor cells within the adult SVZ. The sequencing of 12,200 GFP+ sorted or unselected SVZ cells from this transgenic mouse model has yielded a uniquely comprehensive and granular view of the SVZ stem cell niche uncovering a previously underappreciated heterogeneity. The resultant transcriptional landscape viewed horizontally across time uncovers novel features responsible for reduced neurogenesis with aging.

## Results

Defined components of the endogenous *Nestin* gene promoter/enhancer fragment (*Nes*) that specify expression in NSC have been identified and widely used to express transgenic proteins in the SVZ region (15, 25–29). To increase the power and versatility for identification, targeting, and isolation of adult NSC, we constructed a new transgene, termed *CGD,* designed to express three proteins from one transcript (<u>C</u>reERT2, histone 2B tagged enhanced <u>G</u>FP, and human diphtheria toxin receptor; Fig. 1A & B). The properties of the *CGD* transgene cassette, including appropriate protein expression, Cre recombinase activity and diphtheria toxin receptor (DTR) function were verified in cell culture prior to development of transgenic mouse lines (Fig. S1A-E). Twenty-nine *CGD* transgenic founder lines were screened to identify a single founder that displayed the desired expression patterns in the embryonic and adult central nervous system stem cell/progenitor compartments without detectable ectopic expression (Fig. 1C & D).

**Fig. 1.**
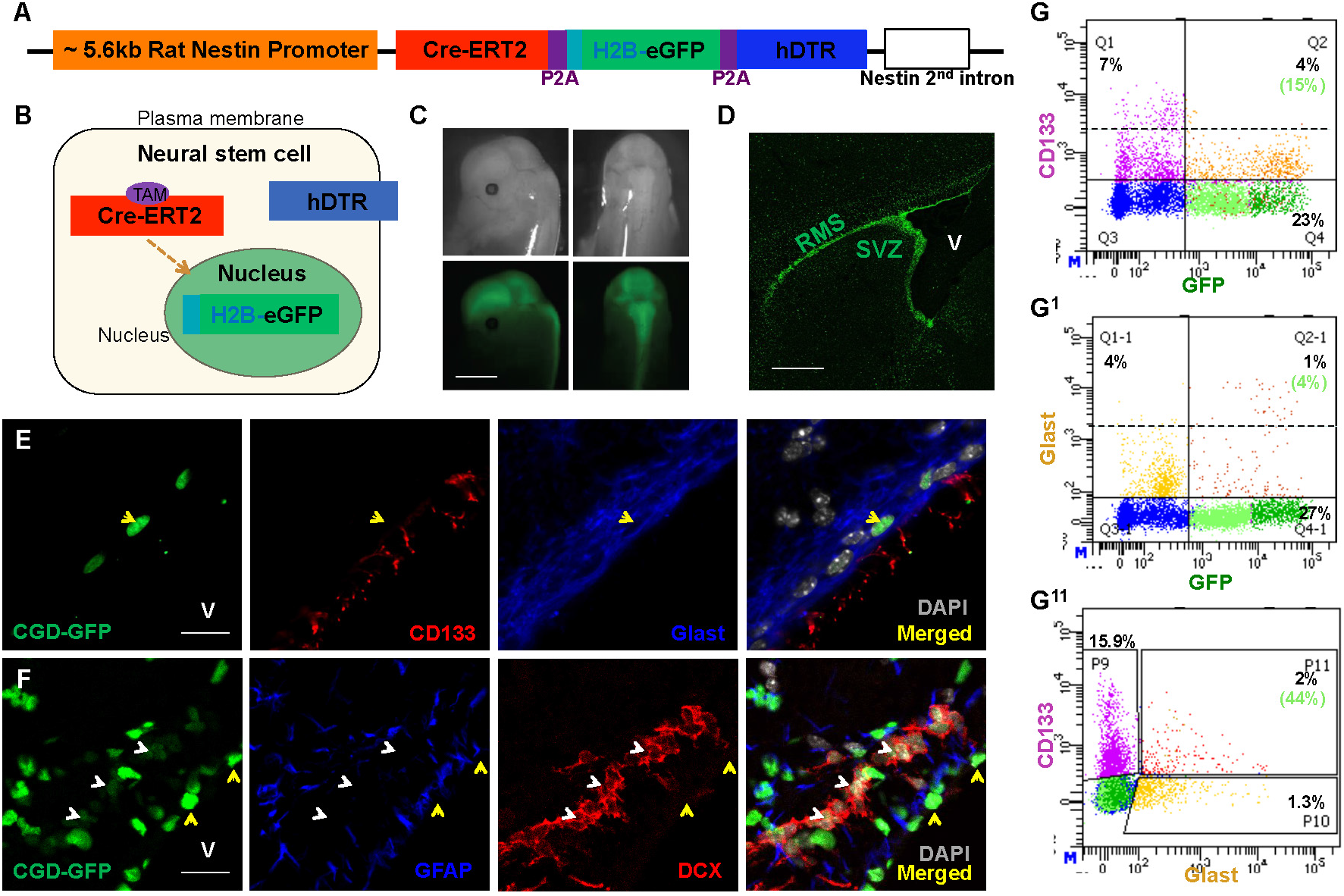
*CGD* transgene labels adult subventricular zone neural stem cells and progenitors. (A) Diagram of the transgene construct. Purple blocks represent the P2A ribosomal skipping element insulating the three gene cassettes. (B) Cartoon illustrates localization of the three transgene products in NSC: Cre-ERT2 in the cytosol, H2B-eGFP in the nucleus, and hDTR on the plasma membrane. (C) Transgene GFP imaging of a 14-day embryo undergoing neurogenesis illustrates *CGD* transgene expression in developing central nervous system. Scale bar: 3mm. (D) Enhanced transgene GFP expression in adult subventricular zone (SVZ) and rostral migratory stream (RMS). Lateral ventricle (V). Scale bar: 500μm. (E) Coronal section staining of two-month old mouse SVZ shows co-localization of *CGD*-GFP with stem cell markers CD133 and Glast. Lateral ventricle (V). Scale bar: 20μm. (F) Relationship between *CGD*-GFP^hi^ and stem cell marker GFAP, versus GFP^lo^ and progenitor marker DCX in a sagittal section of adult SVZ. Scale bar: 20μm. (G-G^11^) A small fraction of *CGD*-GFP+ cells in the SVZ express CD133 (G) or Glast (G^1^), and 44% of all CD133+Glast+ double positive cells express *CGD*-GFP (G^11^). V: lateral ventricle in (D-F). See also Figure S1.

### Functional correlation of GFP+ cells with NSC/progenitor markers

Commonly used markers to identify and label the NSC lineage include the proteins GFAP, Glast, and CD133 (10, 19–21, 23, 24, 30–32). We investigated the expression of these markers in relationship to the *CGD* transgene nuclear GFP reporter. Immunohistochemical (IHC) staining of sagittal and coronal adult brain sections from *CGD* mice confirmed that Glast, CD133, and GFAP antibodies decorated the most intensive GFP+ nuclei (Fig. 1E & F) and wholemount staining of SVZ tissue identified the “pin wheel” structure formed by the GFP+/CD133+/Glast+ NSC surrounded by ependymal GFP negative cells (Fig. S1F) (33). FACS analysis of dissociated SVZ tissue indicated that only a small fraction of *CGD*-GFP+ cells co-express CD133+ (15%), which coincides with the CD133^lo^ category (Fig. 1G, Quadrant 2). Similar analysis of GFP/Glast coexpression indicated that only 4% of GFP+ cells were double positive for Glast (Fig. 1G^1^, Quad. 2-1). Taking into account all CD133+/Glast+ SVZ cells (about 2% of the entire cell preparation), less than half were *CGD*-GFP+. Thus, in the SVZ, *CGD*-GFP reporter expression marks a comprehensive cohort of cells including a significant proportion of CD133 and Glast expressing cells (Fig. 1G^11^, P11). We next examined the relationship of *CGD*-GFP expression to that of doublecortin (DCX) protein, a progenitor marker, and found it excluded from the intensely fluorescing GFP^hi^ cells but instead decorated a cohort of less intense, GFP^lo^, fluorescent cells (Fig. 1F).

While IHC analysis provides a qualitative snapshot of *CGD* transgene expression in NSC/progenitor cells, FACS analysis allows quantification for the whole SVZ tissue. Consistent with the IHC studies, FACS sorting for GFP revealed three distinct populations: GFP^hi^, GFP^lo^, and GFP-cells (Fig. 2A). GFP^hi^ sorted SVZ neurospheres were cultured in defined serum free medium with supplemental growth factors for six days and re-sorted for GFP. A large fraction (~50%) of GFP^hi^ cells transitioned to GFP^lo^ and GFP-states (Fig. 2 A^1^ & A^11^). These data are consistent with a lineage relationship among the varying GFP levels. We next more rigorously tested the self-renewing properties of *CGD*-GFP expressing stem/progenitor cells by evaluating low-density doublet formation and progression to neurospheres. Two hundred dissociated GFP^hi^, GFP^lo^, or GFP-cells were plated per well, in 96 well plates. All wells were examined after sixteen to twenty-four hours for the presence of doublets and again at six days when neurosphere numbers were determined (Fig. 2B-D). Within the first 24 hours, the GFP^lo^ cultures had the greatest proficiency for generating doublets. In contrast the GFP^hi^ and GFP-cells exhibited reduced capacity to form doublets. However, the situation shifted over the subsequent five-day period. The greatest number of neurospheres arose among the GFP^hi^ population (650) whereas the GFP^lo^ doublets yielded fewer neurospheres (250). Thus, despite inefficient initial cell cycle entry and doublet formation, the GFP^hi^ SVZ population shows the greatest capacity to generate neurospheres while the GFP^lo^ population has limited potential to endure in primary culture and form spheres. In this assay, *CGD* transgene expression (GFP^hi^ plus GFP^lo^ cells) accounts for essentially all (99.6%) of primary neurospheres formed in these assays (Fig. 2D). In these studies, GFP-SVZ cells failed to form spheres and to thrive in serum-free medium (Fig. 2D-D^1^). Time-lapse movies of freshly sorted *CGD*-GFP^hi^ cells further revealed an inherent functional heterogeneity among the GFP^hi^ cells. One group of cells formed doublets within 24 hours while a second group remained as singlets for two days or more before entering cell division (Supplemental Movie 1). These data indicate additional functional heterogeneity within the GFP^hi^ cohort of SVZ cells.

**Fig. 2.**
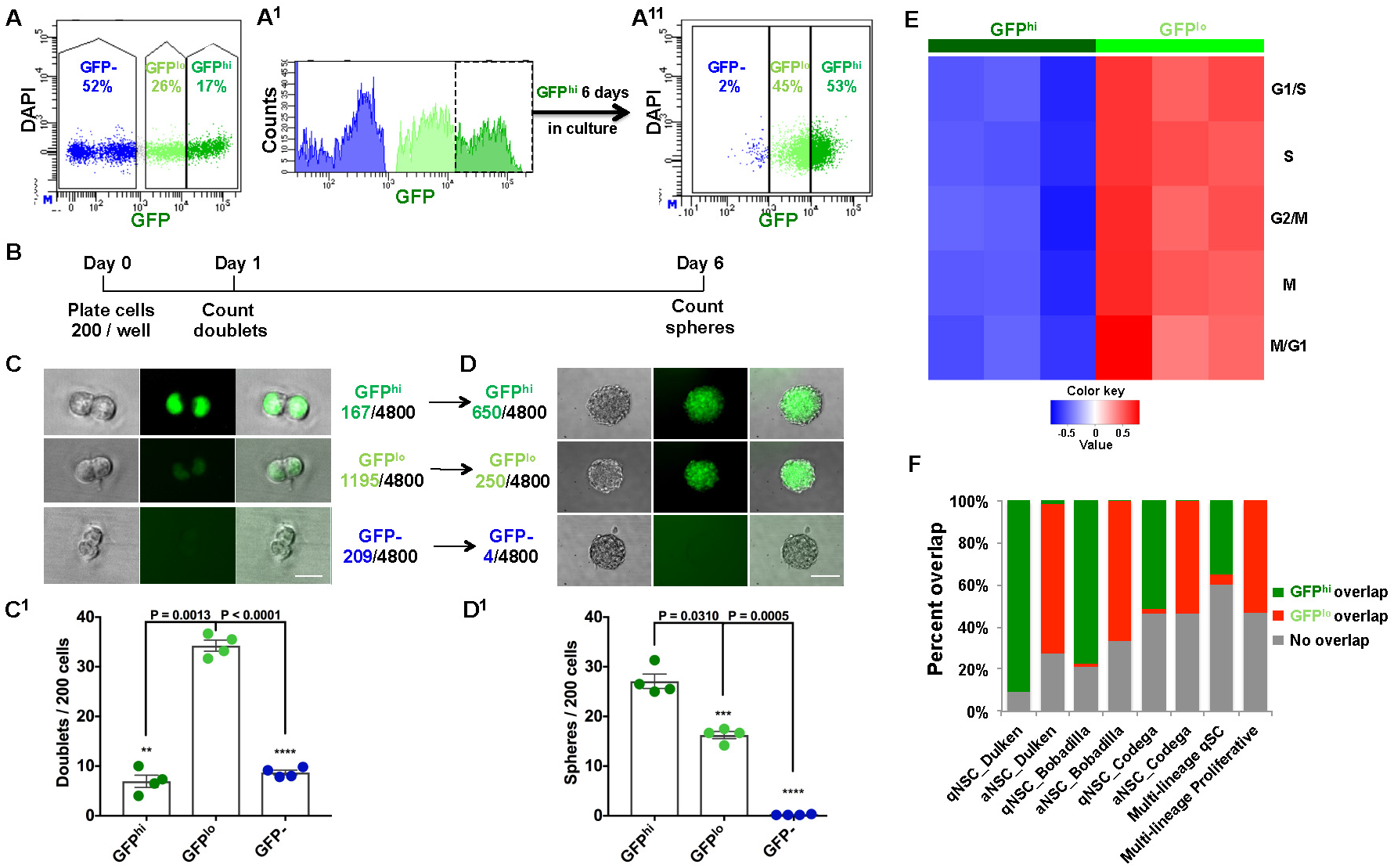
Functional assays validate *CGD* expression in adult neural stem and progenitor cells. (A-A^11^) FACS analysis of whole adult SVZ delineates two GFP populations: GFP^hi^ and GFP^lo^ and GFP-cells. (A^11^) When placed in serum free culture, GFP^hi^ cells sequentially progress to GFP^lo^ and GFP-state. (B) Diagram of timeline for doublet and sphere formation assays below. (C-D^1^) Representative images and statistical analysis of (C-C^1^) 24 hour doublet and subsequent (D-D^1^) 6 day sphere formation assays for FACS sorted adult SVZ GFP^hi^, GFP^lo^, and GFP-cells. Note that despite inefficient doublet formation, the more initially quiescent GFP^hi^ cells are most efficient in forming neurospheres. Mean ± SEM, n = 4 biological replicate mice for each group (each representing 6 technical replicate wells). Scale bars in C and D: 10μm and 100μm. (E) Following RNAseq of GFP^hi^ and GFP^lo^ sorted cells, gene set variation analysis (GSVA) using cell cycle signatures indicates a low proliferative status for GFP^hI^ cells compared to GFP^lo^ cells. (F) Differentially expressed genes (DEGs) of GFP^hi^ and GFP^lo^ cells have high coincidence with published quiescent-stem and activated-progenitor cell signatures. qSC is an acronym for quiescent stem cell (multi-lineage). See also Figure S2 and Table S1.

*CGD-*GFP FACS analysis of dissociated SVZ tissue identifies about 43% of sorted cells as GFP+, with a GFP^hi^ composition of approximately 17% and a GFP^lo^ composition of about 26%. These data coincide with anatomical estimates of cell types present in the adult murine SVZ reporting that Type B cells (stem cells) represent over 20% of the regional cell population (34). In contrast, other SVZ niche studies made use of combinations of markers and/or reporter transgenes to enrich for NSC or progenitors. These different combinations of markers including Glast, CD133, and EGFR, and reporter transgenes like *GFAP-GFP* were reported to yield putative NSC populations that range from 2% to 4.4% of the SVZ preparation indicating a significant underrepresentation of the entire SVZ population (21–23).

To further investigate the *in vivo* relatively quiescent versus proliferative state, freshly dissected and sorted SVZ GFP^hi^ and GFP^lo^ cells underwent bulk RNAseq transcriptome analysis. Gene set variation analysis (GSVA) using five sets of cell cycle phase gene signatures indicated that while the GFP^hi^ cells uniformly exhibit low cell cycle gene expression, GFP^lo^ cells have robust expression (Fig. 2E) (35–37). In aggregate these observations suggest a greater functional complexity among *CGD*-GFP transgene expressing cells in the SVZ that goes beyond the discriminatory powers of established stem versus progenitor protein marker-based approaches.

We also performed gene set variation analysis (GSVA) of the *CGD*-GFP^hi^ and GFP^lo^ bulk sequencing data using published SVZ stem/progenitor cell signatures (21, 23). The differentially expressed genes (DEGs) demonstrated substantial overlap between GFP^hi^ and “quiescent” NSC populations, as did the GFP^lo^ and “activated” NSC populations (Fig. 2F; and S2A-S2B^1^). Gene ontology (GO, https://david-d.ncifcrf.gov/home.jsp) (38) analysis of the GFP^hi^/GFP^lo^ DEGs identified several stem cell signaling pathways enriched in GFP^hi^ cells, including glucose metabolism, lipid metabolic pathway, and oxidation reduction. The GFP^lo^ cells presented cell cycle related signatures including transcription, translation, ribosome biogenesis, etc. (Fig. S2C-E and Table S1C&D). In sum, these results indicate that the *CGD* transgene provides an effective tool to identify and efficiently enrich for SVZ stem and progenitor cells in a comprehensive manner.

### Discrimination of cell subgroups among SVZ NSC/progenitors

To examine whether *CGD*-GFP sorting introduces undue bias in representation of the scope of NSC/progenitor cells in SVZ tissue, we turned to single cell RNA sequencing (see Experimental Procedures). Whole *CGD* transgenic SVZ from three adult mice was dissected, submitted to FACS for cell viability (DAPI-), and approximately 2000 DAPI-cells underwent single cell RNA sequencing. This preparation contained a presumably unbiased mixture of GFP^hi^, GFP^lo^, and GFP-cells (termed UNB). In a parallel experiment, SVZs from six *CGD* transgenic mice were dissected, intermixed and sorted for relative GFP levels (high, low or negative). The Seurat package (http://satijalab.org/seurat/get_started.html) was utilized to process the resulting transcriptomes and cell groupings for both experiments were identified through clustering and visualized using tSNE (t-Distributed Stochastic Neighbor Embedding) projection. We then combined the entire single cell sequencing data collected from the four samples (1,600 GFP^hi^ cells; 1,900 GFP^lo^ cells; 1,100 GFP-cells; and 2,000 UNB cells), resulting in fourteen coherent groupings (Fig. 3A-B). The GFP^hi^ cells dominated four groups (H0-H3); GFP^lo^ cells were enriched in three separate groups (L0-L2); and GFP-cells were ascribed to the remaining seven groups (GFP-;N1-N7; Fig. S3A). The *CGD*-GFP+ (GFP+:H0-L2) cells dominated the stem/progenitor cell lineage (Fig. 3A-B), which accounted for 99% of GFP^hi^ and 92% of GFP^lo^ cells in the SVZ (Fig. S3A). Conversely, GFP^hi^ and GFP^lo^ cells together represent 96% of the cells in subgroups GFP:H0-L2 (Fig. S3A). We independently probed the UNB single cell data set to examine *CGD* transgene transcript expression. *CGD* transcripts could only be appreciably detected in seven subgroups that corresponded to the GFP+ subgroups (GFP:H0-H3 and GFP:L0-L2) ascribed to the sorted GFP^hi^ and GFP^lo^ single cell sequencing data sets (Fig. S3B). Thus, single cell sequencing reveals underlying molecular heterogeneity both within the quiescent (GFP^hi^) and progenitor (GFP^lo^) cell groups that were previously masked in the bulk RNA sequencing of the samples depicted in Figure 2E. We conclude that the *CGD* transgene exclusively and efficiently labels the majority of the SVZ-derived stem cell/progenitor cell pool.

**Fig. 3.**
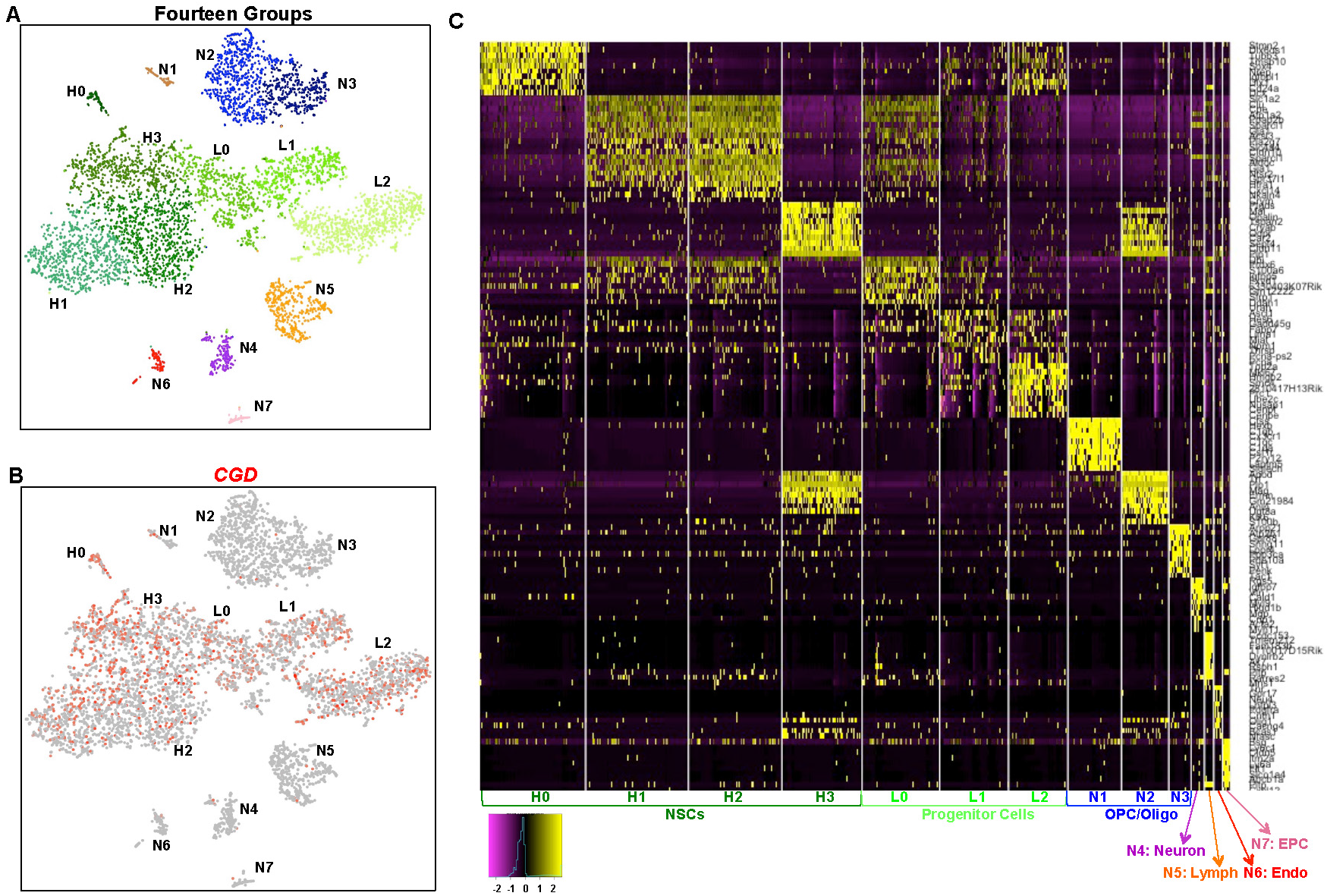
Adult SVZ single cell RNA sequencing analysis reveals seven distinguishable *CGD*-GFP+ groups. (A) tSNE projection of Seurat analysis from the sum of the sorted SVZ samples (UNB, GFP^hi^, GFP^lo^, and GFP- cells) reveals fourteen populations. (B) Normalized *CGD* expression visualized on tSNE coordinates of SVZ cells demonstrates its preferential expression in stem-progenitor populations. (C) DEGs distinguish fourteen SVZ cell types. Heatmap is generated using the top ten unique genes for each group. EPC is an acronym for endothelial precursor cells, Endo signifies endothelial cells, and Lymph signifies lymphocytes. See also Figure S3 and Table S2.

We further analyzed the transcriptomes of the fourteen SVZ subgroups to identify both DEGs and uniquely expressed genes (UEGs) in each subpopulation. The data revealed varying numbers of DEGs, ranging from 114 to 1454 genes, and the UEGs per subgroup ranged from 1 to 866 (Fig. 3C & Table S2 B&C). Not surprising, the GFP-cells are the most heterogeneous (Table S2B) with each of the seven subgroups exhibiting diverse specialized genes and transcriptional pathways representative of the spectrum of cells present in the normal brain parenchyma including: myelination (oligodendrocytes-oligodendrocyte precursor cells; N1-N3), synaptic transmission (neurons; N4), immune regulatory pathways (lymphocytes; N5), and vasculature (endothelial cells and endothelial progenitors; N6-N7), respectively.

Among the GFP+ subgroups, of the classic NSC marker genes, only GFAP mRNA is truly confined to the GFP^hi^:H0-H3 groups although it does not seem to discriminate between them (Fig. S3C). CD133 mRNA showed persistent but low representation in all seven GFP+ subgroups and also in the GFP-:N7 endothelial precursor subgroup (Fig. S3C). Glast transcripts are highest in the GFP^hi^:H0-H3 groups but expression persists throughout the SVZ (Fig. S3C). Similarly, all four GFP^hi^ subpopulations express NSC associated transcription factors including Sox2, Sox9, and ID4 (Fig. 4A, S3C^1^). In contrast, transcription factors associated with progenitor cells such as Ascl1, Dlx1, and Dcx were absent in the four H0-H3 groups, but were enriched in the GFP^lo^ populations (Fig. 4B). The cell cycle gene signatures used to identify the proliferative activity of GFP^lo^ versus GFP^hi^ SVZ cells (Fig. 2E) were applied to the single cell sequencing data and uncovered greater resolution (Fig. S4A). Among the three GFP^lo^ groups, L1 is clearly the most mitotically active and this activity is not uniform with the L0 and L2 groups as implied by the bulk sequencing data. In addition, the GFP^hi^ group data reveals heterogeneity despite the overall limited cell cycle gene expression. The GFP^hi^:H2 group also shows activity in the G1/S phase, which could represent self-renewal (Fig. S4A). GO analysis (Table S2D) associated the four GFP^hi^ subgroups with stem cell related functions including cilium morphogenesis, oxidation-reduction, homeostatic processes, generation of precursor metabolites, etc. GFP^lo^ cells are associated with cell cycle and proliferation genes as well as with signatures of progenitor cells. Thus, single cell sequencing is consistent with the bulk sequencing data for GFP^hi^ and GFP^lo^ cell populations but additionally uncovers details of both the quiescent and progenitor NSC states within the SVZ.

**Fig. 4.**
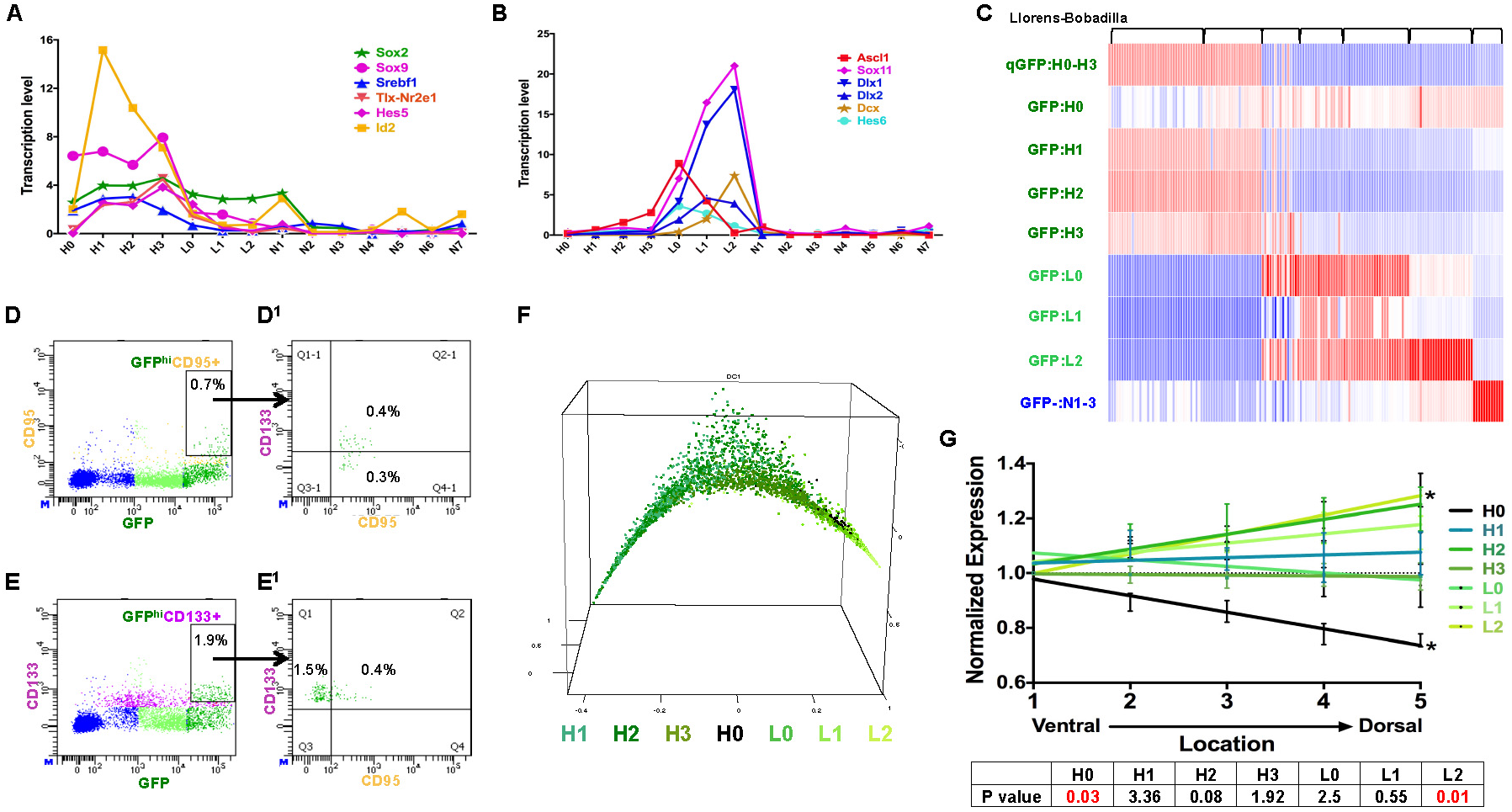
GFP^hi^ cells resolve into four subgroups of quiescent neural stem cells. (A) NSC specific transcription factors are enriched in the GFP^hi^ subgroups (all standard errors < 0.6, units are normalized values scaled to 10,000 counts/cell). (B) Neural progenitor specific transcription factors are enriched in GFP^lo^ subgroups (all standard errors < 0.2, units are normalized values scaled to 10,000 counts/cell). (C) GFP^hi^ and GFP^lo^ subgroup specific gene signatures improve resolution of published SVZ cell sequencing profiles. Transcriptomes of 155 cells published in Llorens-Bobadilla et al., 2015 were analyzed with our signatures. Each column represents one cell. (D) Approximately one half of GFP^hi^;CD95+ cells are also CD133+. (E) About 20% of the GFP^hi^;CD133+ cells are CD95+. (F) Pseudotime analysis provides a nearest neighbor random walk of single cell transcriptional profiles revealing a linear progression from GFP^hi^:H1 through GFP^lo^:L2. (G) H0 cells preferentially reside on the ventral aspect of adult SVZ as defined by Allen Brain Atlas expression profiles. See also Figure S4.

### Four subgroups of quiescent NSC

To identify a core set of GFP^hi^ NSC genes, we compared expression of each of the four GFP^hi^ populations with the remaining subgroups. This yielded 121 commonly expressed genes among the four *CGD*-GFP^hi^ groups (Table S2G). A previous and related study (23) reported RNA sequencing of 130 adult murine SVZ single cells that were presorted for Glast+/CD133+, and further evaluated by EGFR levels (termed qNSC or aNSC respectively). The study also presorted for neuroblasts using a PSA-NCAM antibody (23). We analyzed this published data set by probing for the presence of the gene signatures derived from the fourteen GFP+ and GFP-subgroups including the *CGD*-GFP^hi^ 121 shared gene signature (Fig. 4C, and Table S2G). Significant correlations were observed between the signatures of the differentiated parenchymal cell populations (e.g., GFP:L2 to NB-neuroblasts; GFP-:N1-3 to Oligodendrocytes, Fig. 4C), but not the more quiescent subgroups. To examine CD133 protein expression in GFP^hi^ populations, we performed FACS analysis using CD133 antibodies together with a GFP^hi^:H0 specific marker, Fas-CD95 (Fig. S4B). The results indicate that only half of GFP^hi^;CD95+ (GFP^hi^:H0) cells co-express CD133 protein (Fig. 4D-D^1^). Conversely among GFP^hi^;CD133+ cells, only 20% of the cells express CD95 (Fig. 4E-E^1^). Thus CD133-based sorting of SVZ would preselect only a subset of GFP:H0 cells plus additional GFP^hi^ cells.

We next collectively examined the four GFP^hi^ groups (H0-H3) for expression of biological processes associated with quiescent NSC maintenance and proliferation. This results in a comprehensive list of 1914 genes with eighteen related biological processes (Fig. S4D), some of which overlap with the bulk GFP^hi^ vs GFP^lo^ analysis. Further analysis of the stringent 121 GFP^hi^ subgroup gene list identifies five transcription factors and networks including Sox9, Srebf1 that potentially regulate qNSC maintenance (Fig. S4E-E^11^ and Table S2H).

### A linear progression of the *CGD*-GFP+ stem/progenitor populations

The preceding data are consistent with a developmental relationship from quiescent (GFP^hi^) to proliferative (GFP^lo^) cells. To further examine this model we applied the “Destiny” package designed by dimensional reduction to generate a transition probability from one cell to another according to a random diffusion model (39). Pseudotime ranking of SVZ *CGD*-GFP+ cells begins with the *CGD*-GFP:H1 subgroup and progresses to the L2 population (Fig. 4F).

Recent anatomical and lineage SVZ studies have reported a previously unappreciated level of regional heterogeneity (4, 40). To evaluate whether the transcriptional heterogeneity observed in this study might be reflected spatially in the SVZ, we examined regional distribution of quiescent and progenitor GFP^hi-lo^ (H0-L2) specific transcripts. Ten UEGs from the *CGD*-GFP:H0-L2 groups were examined in the Allen Brain Atlas in situ hybridization brain panel collection (http://mouse.brain-map.org) to assess their expression profile within SVZ. Using brain sagittal views, we partitioned the frontal wall of the SVZ into five equivalent regions with most ventral assigned a value of 1 and most dorsal assigned a value of 5. We found that H0 cells primarily reside in the ventral most SVZ, while the L2 subgroup was enriched in the dorsal region (Fig. 4G).

Anatomical studies have pointed to the “Type B cell” in the SVZ as the adult NSC (7, 21–23, 41). Reports have also pointed to ependymal cells as the source of NSC (8, 10, 42). We examined the “ependymal NSC” specific “hub” genes described by Luo et al., and found them to be specifically enriched in the GFP-:N7 population, which our analysis identifies as endothelial progenitor cells unrelated to the GFP+ NSC/progenitor cells (Fig. S5A-B; Table S2D) (10). GFP-:N7 cells do not have neural stem/progenitor cell properties in culture (Fig. 2D) and the Luo et al. gene set is not present in GFP^hi^ (H0-H3) or GFP^lo^ (L1-L3) cells (Fig. S5A) (10). Thus our analysis excludes the Luo et al., ependymal cell signatures from the SVZ stem cell GFP^hi^ cell cohort.

### GFP:H0 cells have unique NSC properties

The presence of cilia in both Type B NSC and ependymal cells is well established (19, 21, 23, 33). Among the GFP+ cells, our transcriptome analysis appoints a cilia signature specifically to the *CGD*-GFP:H0 subpopulation and with the exception of CD133, absent from the GFP-:N7 subgroup (Fig. S5C and Table S2D). To further characterize the GFP:H0 subgroup, we confirmed its unique gene list with qRT-PCR using SVZ cells that were sorted for single cell sequencing. UNB, GFP^hi^, GFP^lo^, and GFP-cells were examined for GFP:H0 unique gene expression. With some variability, all house-keeping genes (Rplp1, Actb, and Hsp90ab1) tested were expressed in all samples (Fig. 5A and data not shown). The four GFP:H0 specific genes, Tmem212, Dynlrb2, Fam183b, and 1110017D15Rik, were enriched in the GFP^hi^ sample (Fig. 5A). The GFP-sample consistently showed lowest transcription levels for all four genes, with Fam183b and 1110017D15Rik below detection. To specifically isolate GFP:H0 cells for functional analysis, the cell surface marker CD95 proved effective for FACS analysis and CD95+/GFP+ sorting yielded a 0.7% population enriched within the GFP^hi^ cells (Fig. 5B-B^1^). This low percentage was concordant with the low representation of GFP:H0 cells in the entire preparation (Fig. S3D). GFP-cells also contained around 1% of CD95+ cells, which were likely to be GFP-:N7 cells (Fig. S5D and see below). Further analysis of concordant CD95+ expression suggested the CD95+ cells trend with higher GFP protein levels, which coincided with the highest transcription level of *CGD* transgene in GFP:H0 cells (Fig. S4B, S5E-F).

**Fig. 5.**
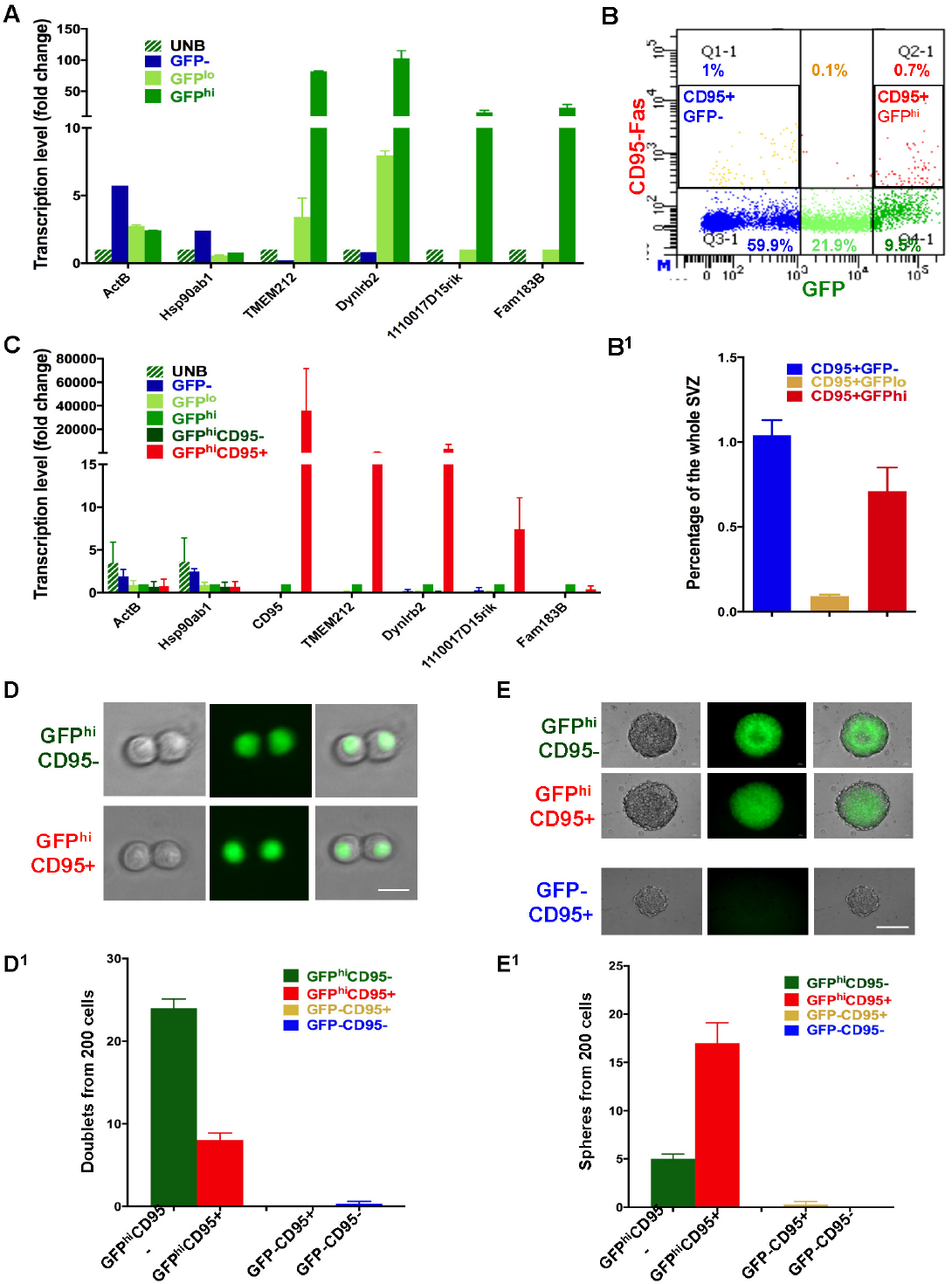
H0 cells have unique stem–like properties. (A) Expression of four H0 specific genes (Tmem212, Dynlrb2, Fam183b, and 1110017D15rik) was verified by Quantitative PCR (qRT-PCR) in single cell sequencing samples. (B-B^1^) CD95-Fas Receptor antibody labels one subgroup of GFP^hi^ and one subgroup of GFP-cells. (C) GFP^hi^;CD95+ cells have enriched expression of the H0 specific genes. qRT-PCR was performed with GFP and CD95 sorted cells for the H0 specific genes. (D-D^1^) Representative images and quantification of doublet formation by GFP and CD95 sorted cells at 24 hours. Note that GFP^hi^;CD95-cells are more efficient at forming doublets. Scale bar: 10μm. (E-E^1^) Representative images and quantification of sphere formation of GFP and CD95 sorted cells. Note that reminiscent of Fig. 2C-D, GFP^hi^;CD95+ cells which were inefficient in doublet formation at 24 hours, are most efficient at forming neurospheres 6 days later. Note also that GFP-;CD95+ cells do not form neurospheres. Scale bar: 100μm. See also Figure S5.

To further assess the identity of *CGD-*GFP^hi^;CD95+ cells as the GFP:H0 subgroup, the four candidate genes used for qRT-PCR validation were reemployed to analyze sorted GFP^hi^;CD95+ and GFP^hi^;CD95-cells. While house-keeping gene expression was present in all samples, the four GFP:H0 specific genes were significantly enriched in GFP^hi^ cells (Fig. 5C). Three were further enriched in GFP^hi^;CD95+ cells (Fig. 5C).

We next compared GFP^hi^;CD95+, GFP^hi^;CD95-, GFP-;CD95+, and GFP-;CD95-cells in doublet and sphere formation assays. Consistent with preceding observations, the two GFP-samples did not form doublets or spheres (Fig. 2C-D and Fig. 5D-E^1^). GFP^hi^;CD95+ cells formed fewer doublets, but more spheres than GFP^hi^;CD95-cells (Fig. 5D-E^1^). These results are compatible with the hypothesis that GFP:H0 (CD95+) cells represent a discrete quiescent NSC population that encounters greater difficulty in entering the cell cycle when initially cultured but outcompetes Type B-like GFP:H1-H3 cells in eventual neurosphere formation. Further, immunostaining of the CD95 antibody with GFP fluorescence confirmed the existence of a double positive cell population residing preferentially in the ventral SVZ (Fig. S5G). Taken together our data indicate that GFP:H0 cells represent a specialized, regionally discrete, quiescent subpopulation of SVZ stem cells.

### *CGD*-GFP+ cell populations decrease with age

We extended the SVZ NSC/progenitor lineage single cell analysis over the process of aging. For this study two-week, one-month, two-month, eight-month, and twelve-month old *CGD* SVZ samples were FACS-analyzed for GFP expression. The results indicated that both GFP^hi^ and GFP^lo^ populations decrease over age. By twelve-months, the GFP^hi^ population was reduced more than nine-fold as compared to the two-week old sample, and the GFP^lo^ population dropped by more than six-fold (Fig. 6A-B). For further analysis, we selected genes with group specific expression profiles in the two-month old SVZ samples (Fig. S6A). Among them, Sox2 was high in all GFP+ cells; E2f1 and Mki67 were specifically induced in GFP:L0-L1; and doublecortin (DCX) was expressed in GFP:L1-L2 subpopulations (Fig. S6A). We found that both GFP+ cells and the L1/L2 markers decreased significantly over age (Fig. 6C-H). While Sox2 decreased only two fold over ten-months, Mki67 and E2f1 decreased by more than four-fold (Fig. S6B). Cells that were double positive for *CGD*-GFP and these aforementioned genes were almost entirely depleted in ten-month samples (Fig. 6C-H & S6C-C^11^). These results indicate that quiescent SVZ NSC gradually lose capacity to enter the cell cycle over the process of aging.

**Fig. 6.**
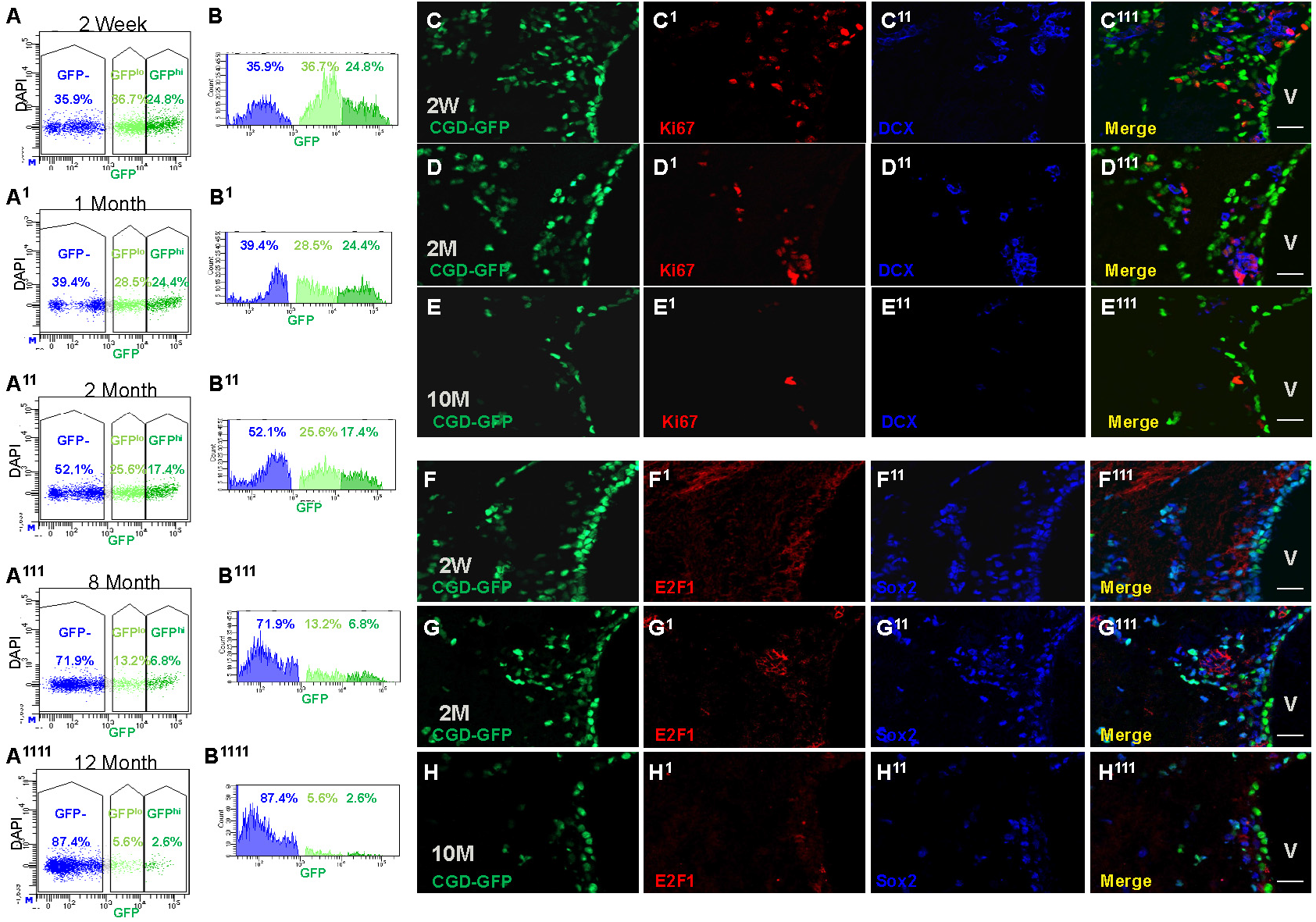
*CGD*-GFP+ cells decrease over age. (A) and (B) Whole SVZ FACS analysis delineates the decrease of *CGD*-GFP+ cells over age. (C) - (E) Commensurate loss of Mki67 and DCX with *CGD*-GFP in two-week (C-C^111^), two-month (D-D^111^), and ten-month mouse brain sagittal sections (E-E^111^). Scale bar: 20μm. (F) - (H) IHC staining for stem/progenitor markers Sox2 and E2f1 with *CGD*-GFP in two-week (F-F^111^), two-month (G-G^111^), and ten-month mouse brains (H-H^111^). Scale bar: 20μm. V: lateral ventricle in (C^111^-H^111^). See also Figure S6.

To examine age related changes for each of the seven *CGD*-GFP+ populations, we performed single cell RNAseq analysis using SVZ tissues acquired at two-weeks, two-months, six-months, and twelve-months. Combined analysis of 5,600 cells identified thirteen major subpopulations (Fig. S7A). The list included thirteen of the fourteen subgroups identified in the original sampling at two months of age, including all seven GFP-subgroups (N1-N7), and six GFP+ subgroups corresponding to GFP:H1-H3 and GFP:L0-L2 (Fig. S7A-D). In this study the GFP:H0 subgroup was underrepresented and did not form a separate cluster but instead merged with GFP:H3 subgroup (Fig. S7A-D).

### Failure of quiescent NSC to transition to progenitors in aged SVZ

We then focused on the GFP+ subgroups and found that two-week SVZ contained the highest proportion of stem/progenitor cells (GFP:H3-L1) concordant with the highest *CGD* transgene expression (Fig. 7A-B). Over a period of twelve months, the GFP:H3-L1 subgroups were increasingly underrepresented (Fig. 7C&D). Doublet/sphere formation assays with GFP^hi^/GFP^lo^/GFP-cells isolated from two-week, two-month, seven-month, and fifteen-month SVZs also indicated a continuous decline in sphere-formation capacity of GFP^hi^ cells during aging (Fig. 7E-F). did not form a separate cluster but instead merged with GFP:H3 subgroup (Fig. S7A-D). Thus, the GFP:H3-L1 bridge, from quiescent to proliferative states, is disrupted indicating that over age there is a deficit in quiescent cell activation.

**Fig. 7.**
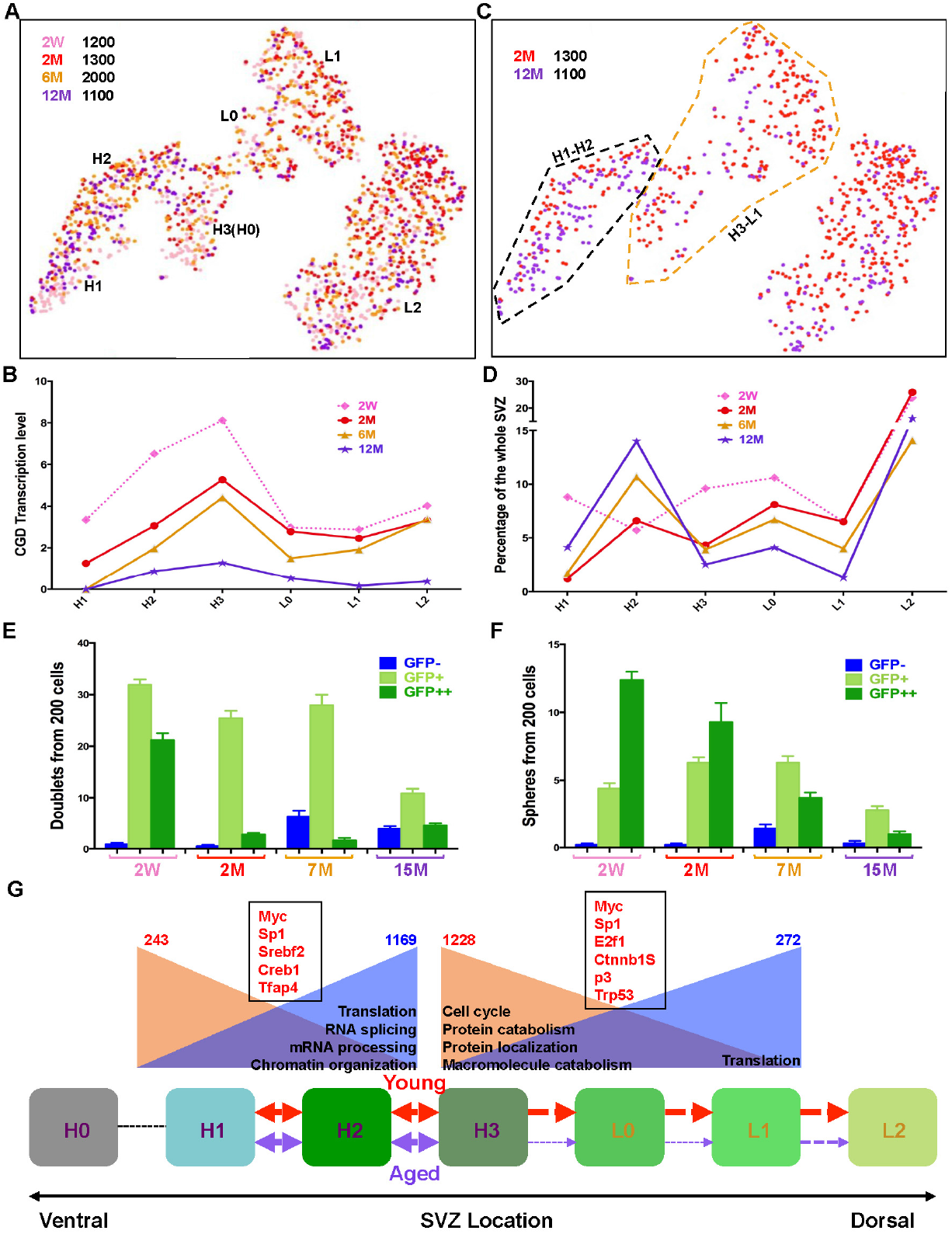
Aging quiescent neural stem cells exhibit deficiency in transition to progenitor state. (A) tSNE projection reveals the neural stem-progenitor lineage distribution of four aged samples: two-week, two-month, six-month, and twelve-month. (B) *CGD* transcription levels consistently decrease over age in each GFP+ subgroup. (C) Direct comparison of the neural stem-progenitor cell distribution between two-month (red) and twelve-month (purple) samples. Note the stark underrepresentation of GFP:H3-L1 lineage in the twelve-month sample. The black dashed line marks the H1-H2 lineage, and the brown dashed line characterizes the H3-L1 cluster. (D) Quantitative illustration of distribution of each GFP+ subgroup (H1-L2) at different age points. Note the continuous decrease of H3-L1 lineage from two-month to twelve-month adult SVZs. (E) and (F) Quantification of doublet and sphere formation assays for SVZ GFP^hi^, GFP^lo^, and GFP-cells over age. Note that by seven months, the GFP^hi^ population begins to lose its capacity to transition from doublets to neurospheres, consistent with the reduced capacity to transition from H3 to L1 in vivo. Mean ± SEM, n = 3 biological replicate mice for each group (each representing 3 technical replicate wells). (G) Cartoon describes the activation blockage of adult NSC in old mice compared to young mice. H1 cells in the SVZ progress into H2 and H3, then become activated to form L0, L1 and L2 cells. This progress is robust in young mice (red arrows), but reduced significantly in aged mice (purple arrows). H1-H2 and H3-L1 cells from two-month and twelve-month are combined to perform differential gene analysis. The numbers on the top reflect the down (red) and up (blue) regulated genes in aged groups. GO analysis from the DEGs unravels potential signaling pathways responsible for the changes. The down (brown background) or up-regulated (blue background) pathways during aging are summarized. The red genes in black boxes reflect the putative transcription factors associated with the age-related gene expression change in either H1-H2 or H3-L1 lineages. See also Figure 4F, S7 and Table S3.

To further examine the transition deficit, we compared pooled SVZ GFP^hi^ subgroups (H1-H2, Fig. 7B, black dash-line) and GFP^hi-lo^ transition subgroup (H3-L1, Fig. 7B, Brown dash-line) between two-months and twelve-months. The twelve-month H1-H2 lineage samples yielded 1169 up and 243 down-regulated genes while the H3-L1 lineage yielded 272 up-regulated genes and 1228 down. GO analysis indicated key features of both aged SVZ cohorts including increased translation. The H1-H2 cluster presented additional activity in RNA splicing and mRNA processing, and chromatin organization processes (Fig.7G and Table S3D-J). The aged mice H3-L1 lineage showed downregulation of cell cycle, protein catabolism and localization, and macromolecule catabolism processes. The Transcription factors responsible for the up and down-regulation of the gene sets in each cluster were also determined. While Myc and Sp1 are related to the aging profile switch in both lineages, Srebf2-Creb1-Tfap4 and E2f1-Ctnnb1-Sp3-Trp53 are associated in either H1-H2 or H3-L1 clusters (Fig.7G). These biological processes and transcription factors provide potential targets to overcome aging deficits of NSC.

## Discussion

Single cell tSNE analysis of the dissected SVZ region identified fourteen distinguishable transcriptional subgroups of cells including seven stem/progenitor subgroups. Among them, four superimposed on the *CGD*-GFP^hi^ subgroups, three on the *CGD*-GFP^lo^ subgroups, and the remaining seven groups on the GFP-subgroups. This provides powerful validation that the *CGD*-GFP sorting process had indeed portrayed an unbiased representation of the stem/progenitor cell niche.

Functional assays including primary culture low-density doublet and neurosphere formation indicated that the sorted GFP^hi^ cells represent a quiescent stem-like cell population. When placed in culture, this population is most efficient in establishing neurosphere cultures when compared to the GFP^lo^ cells that efficiently form doublets but have reduced potential to evolve into neurospheres. Neurosphere formation is widely considered as a valid assay for stem cell self-renewal (43–47). However, as we show, transient amplifying cells can also form and sustain neurospheres. Therefore, assays such as the low-density doublet assay and subsequent quantification of progression to neurospheres may provide a more stringent method for assessing self-renewal capability.

A key determinant that permitted analysis of single cell sequence data from different experiments for each of the fourteen SVZ subgroups was the consistent expression of cell specific transcription factors and markers commonly used to define stem cells, progenitor cells, and specialized parenchymal cells (see Fig. S3). Moreover, molecular analysis further validated the identity of GFP^hi^ and GFP^lo^ cells as stem and progenitor cells respectively. GO and cell cycle analysis confirmed the differential status of these cells both in the metabolic and in the cell cycle states (Fig.S4D and Table S2). In contrast the seven GFP negative (GFP-:N1-N7) cell subgroups aligned with parenchymal cell types included in the SVZ preparation. The increased granularity achieved with sequencing of many thousand cells enriched with a novel transgene revealed unexpected features. For example, only two of three proliferative GFP^lo^ subgroups (GFP:L0 & L1) are highly active in the cell cycle. Moreover few of the standard markers for quiescent stem cells segregated at the transcriptional level to a specific subgroup.

The GFP:H0 subpopulation is particularly intriguing and has unique features. Anatomically, GFP:H0 genes preferentially localize in the ventral SVZ. This subgroup is the only one amongst GFP+ cells that has high transcription of cilia genes. Furthermore, isolation of GFP:H0 cells reveals that they are likely the subgroup within the GFP+ cells responsible for reduced low-density doublet formation but enhanced neurosphere development in primary culture. We suggest that H0 cells may represent the most quiescent NSC cell population and whether it participates actively in the homeostatic process of self-renewal and production of transient amplifying cells merits further investigation.

The uncovering of additional subgroups of GFP^hi^ cells and of GFP^lo^ subgroups underscores the heterogeneity underlying a complex biology. The identification of subgroup specific transcriptional signatures and importantly, of transmembrane receptor encoding genes should provide the means to prospectively isolate these populations and better study their functions. Very recent studies have used related approaches to investigate the SVZ and our present data serve to validate but importantly to greatly enrich and resolve the emerging picture of the heterogeneous SVZ. We cannot ignore a recent study that through sequencing of twenty-eight SVZ cells, reported a NSC specific signature and proposed an ependymal origin for NSC (Luo et al., 2015). Our sequencing studies, which included six thousand-six hundred cells, ascribes the Luo et al., NSC signature, not to NSC, but rather to endothelial progenitor cells (GFP-:N7 subgroup). Finally, unlike the relative facility in culturing doublets or neurospheres from GFP+ cultures (albeit with differing efficiencies), we were unable to obtain cultures from GFP-cells. This substantial discrepancy raises a cautionary note about the importance of validating single cell data not only at the molecular level but also with meaningful functional assays.

It has long been appreciated that over age, SVZ neurogenesis declines (48–50). However, the effect of aging on different NSC/progenitor populations is unresolved, in part due to methodological constraints of cell identification criteria or limited cell number (49, 51). Our approach allowed detailed transcriptional examination of the process and demonstrate persistence of two qNSC populations (H1 and H2) from two to twelve months. Instead, a particular bottleneck within the stem-progenitor lineage at the H3-L1 juncture becomes pronounced with age. The data provide a rich list of genes including transcription factors and signaling pathways that may be valuable in identifying potential molecular determinants of this deficiency and potential strategies to overcome it. Here, we focused analysis between adult SVZ of two-month and twelve-month. Additional comparisons focusing on the adolescent (two-week) and aged (twelve-month) data may reveal additional insights in NSC maturing and aging. Further analysis using the seven GFP- populations could identify aging related changes in oligodendrocytes, neurons, immune cells, and endothelial cells.

NSC have important functions in normal physiology that warrant their thorough study and understanding. Furthermore, NSC may be among the few or unique cells in the several hundred million cells within the adult brain that have capacity to engender malignant brain tumors such as glioblastoma (18). This disease stands out as one of the most intractable. In this study, we present seven stem/progenitor populations to lay the foundation for further investigation into the mechanism underlying the transition from a stem cell to a tumor cell of origin.

## Materials and Methods

### Animal Studies

All mouse experiments were approved and performed in accordance with the guidelines of the Institutional Animal Care and Research Advisory Committee at the University of Texas Southwestern Medical Center and the Institutional Animal Care and Use Committee of Memorial Sloan Kettering Cancer Center.

### Generation of the *CGD* transgenic mice

A *CGD* fragment was amplified and assembled from a CreERT2 containing construct, an H2B-eGFP vector (from Yan Jiang, Mount Sinai), and genomic DNA from 293T cells (diphtheria toxin receptor). To ensure the efficient expression of all the three proteins in the *CGD* transgene, a P2A fragment was used to space between the three different genes (52–54). To verify the adequate expression of mono-proteins from the multi-cistronic cassette, the coding region of the *CGD* fragment was inserted into a FETPTR vector to transfect 293T cells. Western blot was used to detect the comparative expression levels of monomers vs polymers from the same open-reading frame (ORF) with a GFP antibody (1:1,000, Fig. S1C). Further analysis demonstrated the activity of the three proteins (Fig. S1B-E). The *CGD* fragment was then inserted into the pNERV vector (gift from Steve Kernie, Columbia University) (28) through NotI (New England Labs) sites. The pNERV-*CGD* plasmid was cut with SalI to release the Nestin-*CGD* fragment to generate transgenic mice by pronuclear injection into fertilized murine eggs in a C57BL/6 genetic background. 82 founders were screened with PCR and 29 lines were identified as positive with the *CGD* transgene. The primers used to verify the transgene were: forward primer, 5’-GTCTATATCATGGCCGACAAGC-3’ and the reverse primer, 5’-GAAAGAGCTTCAGCACCACC-3’.

### Histology and Immunohistochemistry

Mice were perfused and fixed in 4% paraformaldehyde, soaked in 30% sucrose, and imbedded in OCT compound (Tissue-Tek). Frozen brain tissues were sectioned (12**μ**m) and stained with antibodies as follows: goat anti-GFAP (Santa Cruz, 1:500), mouse anti-GFAP (1:200, Millipore), goat anti-Doublecortin (Santa Cruz, 1:500), goat anti-Sox2 (Santa Cruz, 1:200), rabbit anti-Mki67 (Novus, 1:500), goat anti-GFP (Rockland Immunochemicals, 1:200), mouse anti-Glast (Miltenyi Biotec, 1:100), rat anti-CD133 (eBioscience, 1:100), rabbit anti-E2f1 (Abnova, 1:100). DAPI was used to stain the nucleus (ThermoScientific, 1ug/ml). The sections were then imaged with a Zeiss LSM 510 confocal microscopy using Argon 488, He543, and He 633. Whole mount SVZs were prepared and stained as described (55).

### Tissue Culture and FACS Analysis

Fresh mouse SVZ tissues were dissected and cultured for doublet and sphere formation analysis with a modified protocol as described (47, 56). Briefly, 200 cells were plated into 96-well plate and counted for doublets within 16-24 hours, and spheres six days later. When cell numbers were limited, 100 - 300 cells were sorted and plated. The data were then normalized to 200 cells. Antibody staining for FACS was performed in Flow Cytometry Staining Buffer (ThermoFischer Scientific) and following antibodies were used: PE-GLAST (ACSA-1, Miltenyi Biotec, 1/40), APC-CD133 (13A4, Miltenyi Biotec, 1/75), APC-Cy7-CD45 (BD, 1/200), APC-CD95-Fas (Miltenyi Biotec, 1:100). DAPI was used to stain the dead cells (ThermoScientific, 1ug/ml). Viable cells were sorted in a FACSAriaII (BD Biosciences) and collected in complete serum-free medium for culture. For the movie in Fig. S2A, the cells were incubated in Cytation 5 (BioTek) and imaged every hour for GFP and cell morphology.

### Total RNA Extraction and Gene Expression Profile Analysis

Freshly prepared samples were sorted directly into concentrated Trizol LS reagent (Invitrogen 10296028) and submitted to Integrated Genomics Operation facility (MSKCC) for RNA extraction and sequencing with paired-end 50 method. The results were processed by Bioinformatics Core in MSKCC to identify differentially expressed genes.

## Acknowledgments

We thank Omar Aly (Weill Cornell Medicine), Yanjiao Li, and Alicia Pedraza for technical support in single cell sample preparation. *H2B-eGFP* plasmid was a gift from Yan Jiang (Mount Sinai) and *Nestin* enhancer/promoter was a gift from Steven Kernie (Columbia University). We thank Elvin Feng (MSKCC Molecular Cytology Core Facility) for designing the ImageJ script used for quantification of gene expression in the Allen Mouse Brain Atlas. We thank Hui Julia Zhao and Nicholas Socci (MSKCC Bioinformatics Core Facility) for bioinformatics support. This work was supported by NIH grants R01 CA131313-01A1, NOA 1R35CA210100-01, and in part through the NIH/NCI Cancer Center Support Grant P30 CA008748. X. P. Xie was supported by American Brain Tumor Association Basic Research Fellowship in honor of Joel A. Gingras, Jr. L.F. Parada holds the Albert C. Foster Chair in Cancer Biology.

## Author Contributions

LFP, XPX, and DRL designed the study and wrote the manuscript together with assistance from OE. DRL, XPX, AP, and AML designed, performed, and analyzed single cell sequencing. XPX designed and performed molecular and cellular experiments with assistance from ZW, GB, DRL, DS and JS.

## Supplementary Information Text

### Supplemental Materials and Methods

#### Single cell sequencing

The Dropseq cell-bead collection, sample preparation, library preparation, and sequencing were performed as previously described (1). Beads were purchased from Chemgenes (#MACOSKO-2011-10) and the microfluidics chip was purchased from Nanoshift. All our reagents were purchased from the recommended sources as outlined in Macosko et al., 2015 and the McCarroll lab’s online Dropseq protocol, v3.1, Dec. 2015 (http://mccarrolllab.com/download/905/). Our primers and oligonucleotides were purchased from IDT with the identical sequences outlined by Macosko et al., 2015. Sequencing was done on a NextSeq 500 at the Cornell Sequencing core, 64bp (R2). Alignment of reads and generation of cellular expression counts were performed with the Drop-seqAlignmentCookbookv1.2Jan2016 (http://mccarrolllab.com/wp-content/uploads/2016/03/Drop-seqAlignmentCookbookv1.2Jan2016.pdf) and with the Dropseq tools provided by the McCarroll lab (http://mccarrolllab.com/download/922/). We utilized the mm10 reference genome into which we incorporated the *CGD* transgene mRNA sequence. Mus_musculus.GRCm38.82 was used as our refflat GTF. The Digital Expression NUM_CORE_BARCODES output was set to 5000 as we knew we had less than or equal to 2000 cells per sample.

#### Single cell analysis by “Seurat”

All our analysis was performed in R version 3.3.0 with R Studio. The “Seurat” R package was used to filter, normalize, cluster and select differentially regulated genes for each cluster. Seurat package (version 1.4.0.16 and, when released, version 2.0) was utilized as described in the online clustering tutorials by the Satija lab (http://satijalab.org/seurat/get_started_v1_4.html, http://satijalab.org/seurat/pbmc3k_tutorial.html) (1). Cells were excluded if genes were detected in less than 3 cells, or had less than 200 unique genes. The resultant matrix was normalized to 10,000 transcripts, log transformed, and a regression to the number of unique molecular identifiers (UMIs) was performed before dimensional reduction. Principle components 1-19 were used to identify subpopulation clusters. We compared the lists of differential genes for each group in a cohort to select genes that were unique to a group, in other words, they were signature genes that were differentially regulated in a certain group but not in any other group. This produced two lists, differentially up-regulated genes and unique up-regulated genes. We used the differentially up-regulated gene list for gene ontology analysis by DAVID (2), (https://david-d.ncifcrf.gov/summary.jsp). We used the top 10 unique genes (DEGs when not enough, Table S2B & 2C) for each group to produce the heatmap with the Seurat package (Figure 3C). We performed batch correction in Seurat (version 2.0) to cluster and project both the Aged samples and the sorted two-month samples together (Fig. S7B-C). To do this we used the union of the top 2000 most variable genes for each data set and ran the canonical correlation analysis (CCA) in Seurat that identifies common sources of variation between the two datasets. For this analysis we used the first 19 dimensions and all the other settings and analyses were the same as above.

When Seurat version 3 was released, we re-did both the sorted wildtype and aged samples analyses with the latest version (data not shown). As the samples were from the same tissue, preparation date, sequencing run, and there were no discernible batch effects in the sorted samples (GFP^hi^, GFP^lo^, GFP-, Unsorted) we did not use the integration method for this analysis but rather simply merged the samples into one matrix before the Seurat pipeline so as not to introduce any noise from unnecessary processing. In contrast, for the aged samples we used the anchor integration method (Stuart et al., 2019). In both instances, the original results were conserved and proved robust across different pipelines and versions of Seurat. One exception was the coclustering of H1 with H2 into one cluster in the Seurat version 3 processing of the sorted two-month SVZ samples. As the results and clustering were largely conserved across different versions of Seurat, we present the original analysis with confidence that they are robust results irrespective of the Seurat versions and processing employed.

#### Pseudotime Analysis

Pseudotime analysis was performed using the Destiny package as detailed in (3) and in the vignette at https://www.helmholtz-muenchen.de/icb/research/groups/quantitative-single-cell-dynamics/software/destiny/index.html. We used H0-L2 groups from Fig. S3A for our Pseudotime analysis in Fig. 4F. In Destiny, we plotted the result with our established color codes. The animation was compiled in Sequimago and labels were added in Wondershare Filmora (Version 7.8.9). Pseudotime analysis was also performed in a similar manner on the H and L groups of the aged samples that were included with the aforementioned pseudotime matrix of H and L groups in Fig. 4F (Fig. 7B).

#### Allen Brain Atlas SVZ localization of unique genes

The gene list used for this experiment is the same as those used in the aforementioned Heatmap section (see above). We collected images from the Allen Brain Atlas (4), http://mouse.brain-map.org/) where the SVZ was at its largest in the sagittal sections. We authored a macro in ImageJ that got rid of the white space in our quantification. We then made 5 sections of the SVZ from Ventral to Dorsal and used ImageJ to quantify the in situ hybridization of RNA for each of the five sections. We then normalized the values to the most ventral section and plotted the values, performed linear regression, and in GraphPad (Prism7) we calculated whether the slopes were significantly different than 0. The p-values for this test are reported.

#### Venn Diagrams

Venn diagrams were created by InteractiVenn (http://www.interactivenn.net/) (5) and reproduced in Photoshop CS6 (Version 13.0.6) to improve the quality and size of the associated text.

#### Transcription factor analysis

Transcription factors related in a sign sensitive manner to our inputted lists of up- and down-regulated genes were identified by TFactS by target gene signatures curated from microarray gene expression data (http://www.tfacts.org) (5, 6).

#### String network

Gene networks were generated by String database (https://string-db.org) (7) with medium confidence (0.400) to assess known and predicted interactions (physical and functional) between genes in our inputted gene sets. The PPI enrichment score p-value is a measure of the significance of enrichment for network interactions within a gene set.

#### qRT-PCR with CD95+ sorted samples

We sorted 1000 GFP^hi^;CD95+, GFP^hi^;CD95-, bulk GFP^hi^, GFP^lo^, GFP-, or DAPI-viable cells into tubes containing 3000 Dropseq beads (Chemgenes) in 100ul lysis buffer with DTT (#15508013, Thermo Scientific) (1). The tubes were rotated at 4 degrees for 10 minutes. Then we washed the samples twice with 6xSSC (1), followed with another wash with reverse transcription buffer (Thermo Scientific). Reverse Transcription mix with Template Switch Oligo were added and incubated for 30 minutes with rotation at room temperature and an additional 90 minutes with shaking at 700 rpm at 42 degrees Celsius. We then processed the samples as Dropseq samples outlined in Macosko et al., 2015 with Exonuclease treatment, PCR for 30-35 cycles, and Agencourt Ampure XP (#A63881, Beckmam-Coulter) bead purification of the resultant DNA. The purified DNA content was measured on a Qubit with high sensitivity reagents (#Q32854, Thermo Scientific). qRT-PCR reactions were performed with SYBR Select Master Mix (#4472903, Applied Biosystems) in an Applied Biosystems QuantStudio Flex6, real time PCR system. The following primers were selected from PrimerBank (8), https://pga.mgh.harvard.edu/primerbank/index.html) except for Cd95 which was selected from Sabbagha et al. (9), and all primers were purchased from Eton Biosciences Inc.

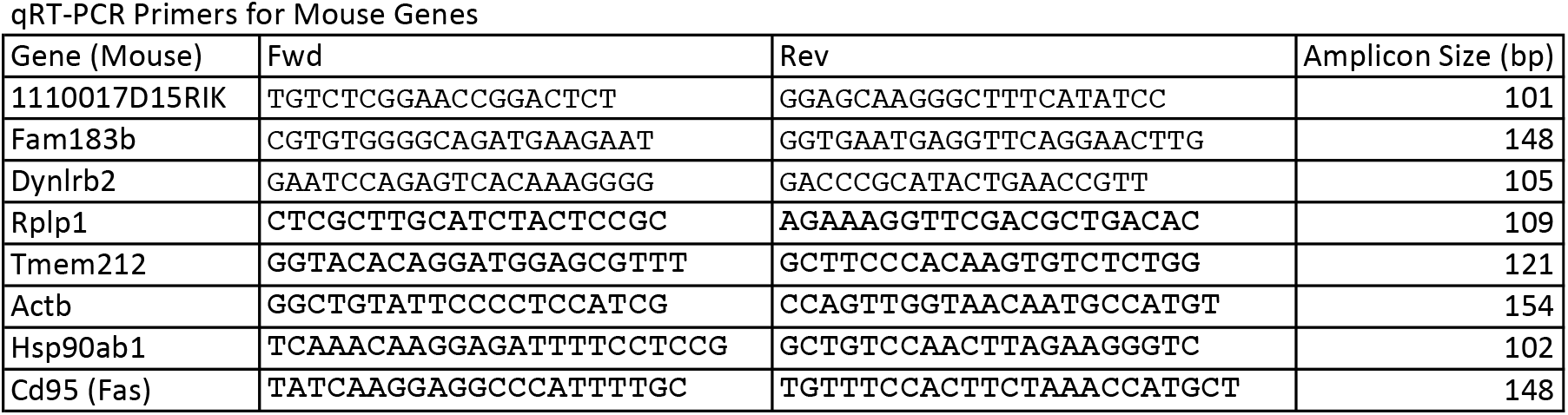

#### SCDE analysis

We employed the package SCDE (https://www.nature.com/articles/nmeth.2967) (10) to determine differentially expressed genes between 2 Months and 12 Months aged samples in both the combined H1-H2 groups (N=93 for 2 months, N=115 for 12 months) and in the combined H3-L0-L1 groups (N=84 for 2 months, N=63 for 12 months). We used non-log transformed, ‘Seurat’ normalized data matrices as the input. We chose genes with FDR P values <0.05 for further analysis including gene ontology (David, https://david-d.ncifcrf.gov/summary.jsp) (2).

**Fig. S1.**
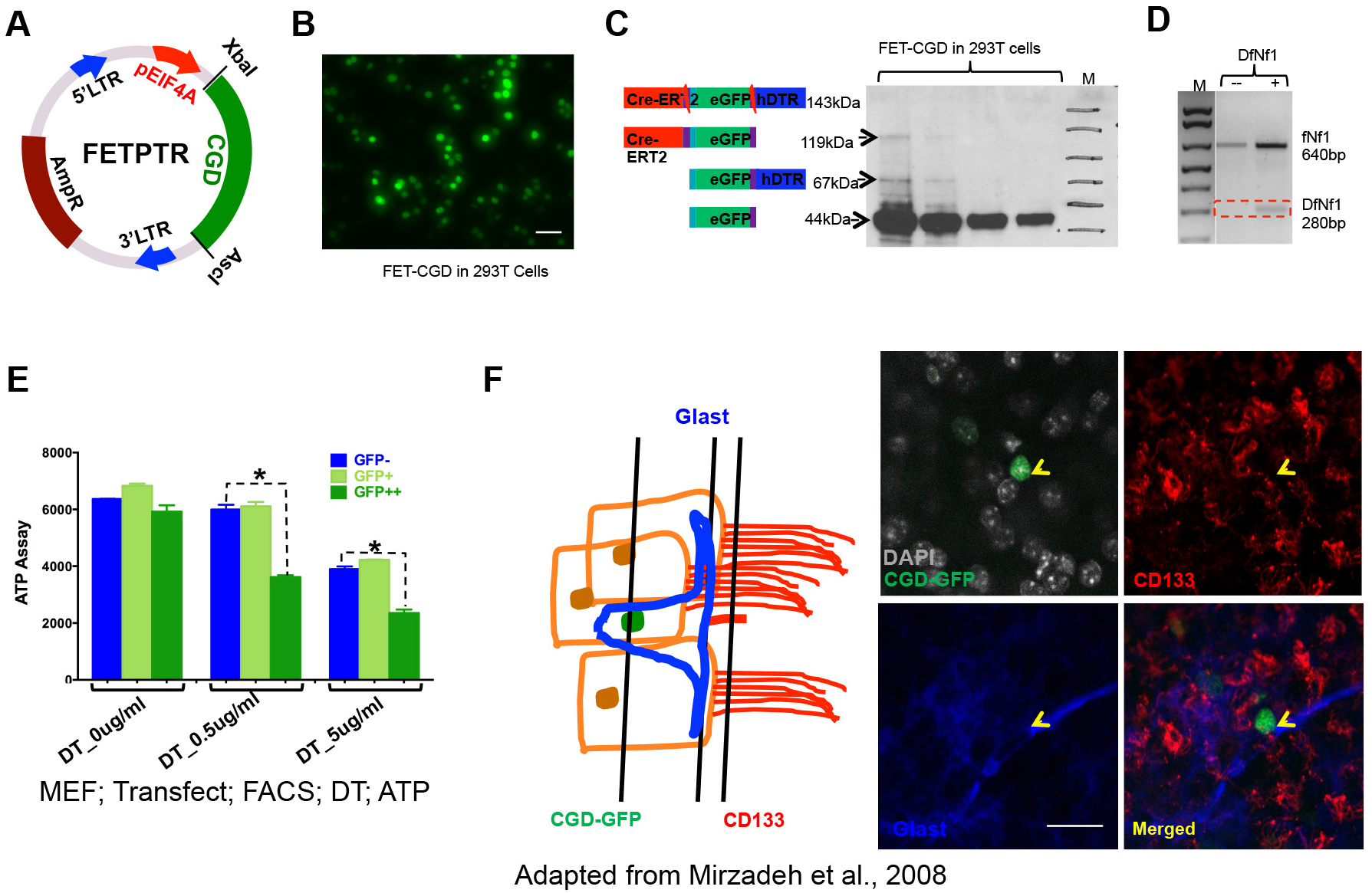
This figure accompanies Figure 1. *CGD* transgene construct characterization. (A) FETPTR expression vector. (B) eGFP reporter detection in FETPTR-*CGD* construct transfected 293T cells. Scale bar: 20μm. (C) Western blot with GFP antibody verifies efficient production of GFP monomers. 293T cells were transiently transfected with virus packaging a pEIF4A promoter driven *CGD* cassette and collected three days later in 2X laemmli for western blot. (D) CreER protein from the *CGD* transgene can remove the floxed region within an NF1 gene allele, detected by PCR. (E) Mouse embryo fibroblasts transfected with the FETPTR-*CGD* derived viruses are sensitive to diphtheria toxin in an ATP assay. (F) Left: Diagram of classic SVZ “pinwheel” structure in which a eGFP+/Glast+/CD133+ stem cell is surrounded by CD133+ ependymal cells. The three black lines indicate the relative positions for the microscope images on the right. Right: Corresponding IHC SVZ images with a *CGD*-GFP^hi^ nucleus and Glast+ cytoplasm surrounded by CD133+;GFP-ependymal cells. Scale bar: 20μm.

**Fig. S2.**
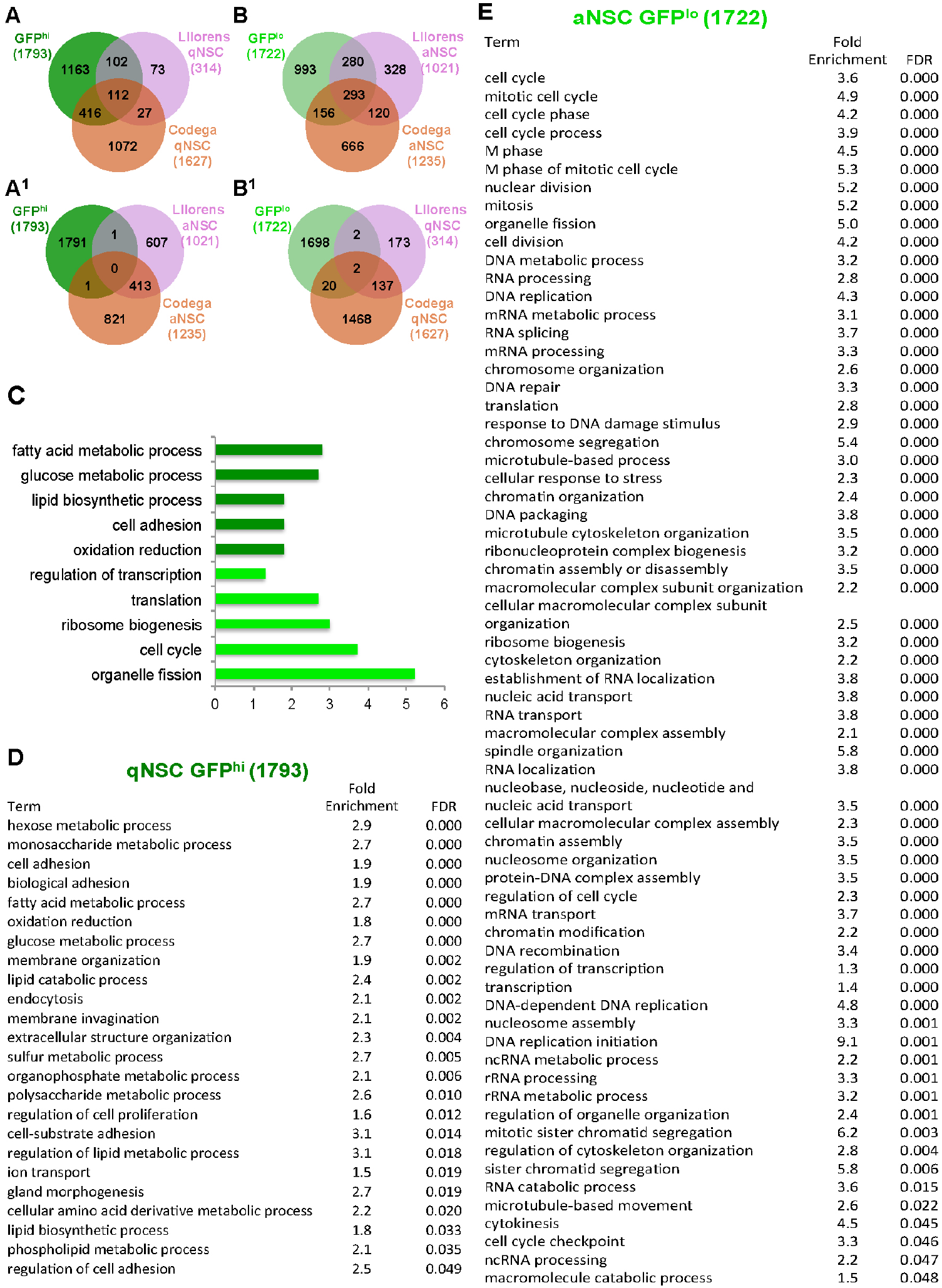
This figure accompanies Figure 2. Gene expression analysis of *CGD*-GFP^hi^ and GFP^b^ cells point to a quiescent versus a proliferative state respectively. (A) Venn diagram analysis comparison of *CGD*-GFP^hi^ DEGs and published quiescent NSC signatures demonstrates significant overlap but not with activated NSCs (A^1^). (B) GFP^lo^ cell signatures preferentially overlap with that of published activated NSCs but not with quiescent NSCs (B^1^). (C) Gene Ontology analysis of *CGD*-GFP^hi^ and GFP^lo^ profiles is consistent with a stem versus progenitor state. (D) and (E) Known and unknown biological processes related to quiescent or proliferative NSCs are associated with GFP^hi^ (D) or GFP^lo^ cells (E).

**Fig. S3.**
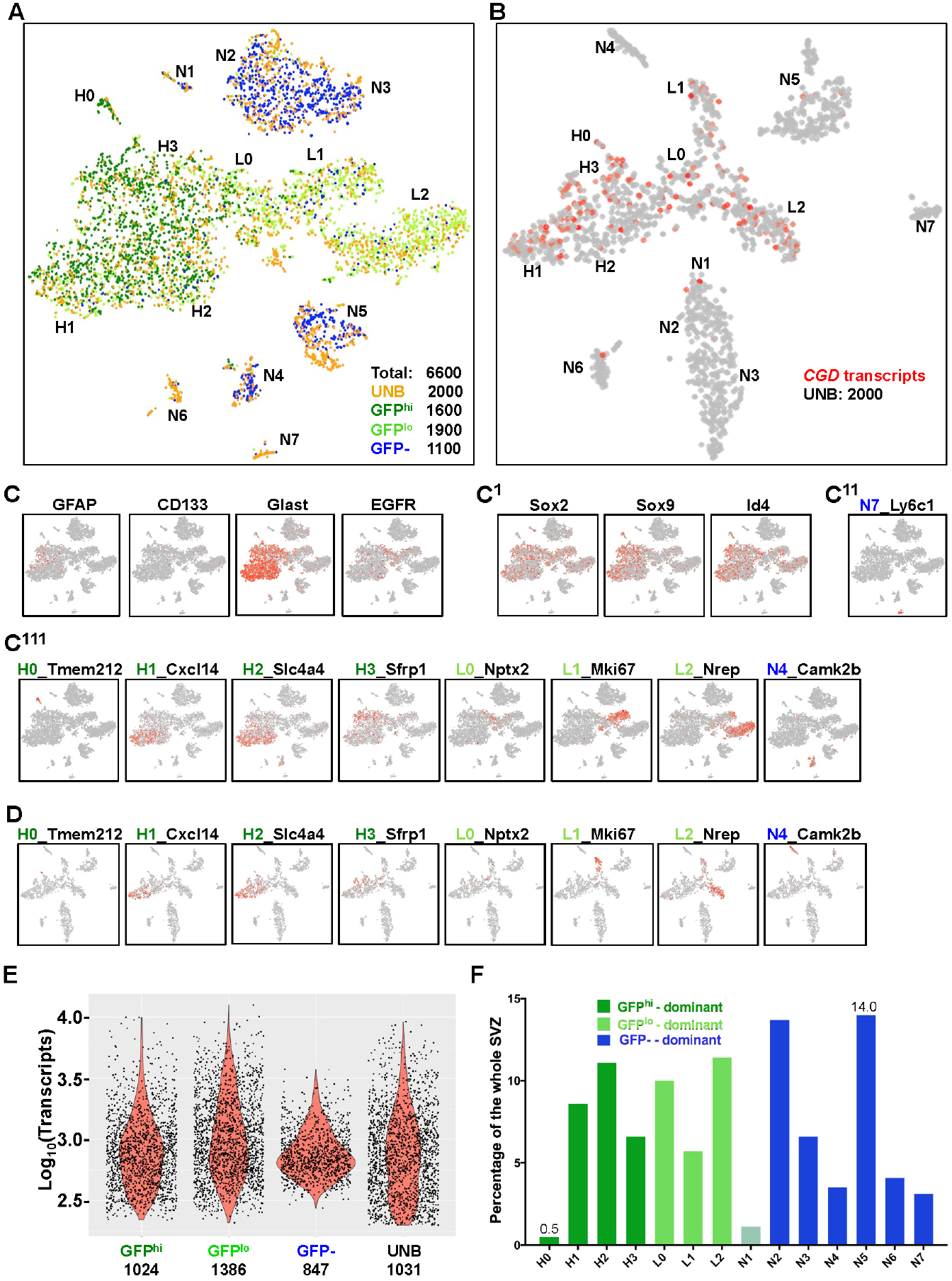
This figure accompanies Figure 3 and Table 2. Drop-seq analysis of whole SVZ. (A) Combined 6600 cells yield fourteen groups that integrate the GFP^hi^, GFP^lo^, and GFP-subgroups. Note UNB cells exist in each of the fourteen groups. (B) tSNE projection of UNB cells shows similar groups and enriched *CGD* expression in H1-L2 lineage. (C) Distribution of sorting markers for stem/progenitor cells used in previous studies. (C^1^) Representative distribution of NSC and progenitor specific transcription factors; (C^11^C^111^) candidate markers identified in this study for GFP-:N7 and different SVZ stem/progenitor groups. (D) Representative markers for NSC/Progenitor/neuron lineage facilitate the identification of respective populations in UNB cells. (E) Transcript numbers for each of the four sorted and Dropseq analyzed samples present a “bell” shape distribution. Average transcripts per cell for each sample are indicated at the bottom. (F) SVZ UNB cells contain fourteen distinct subgroups ranging from 0.5% to 14% of the whole tissue.

**Fig. S4.**
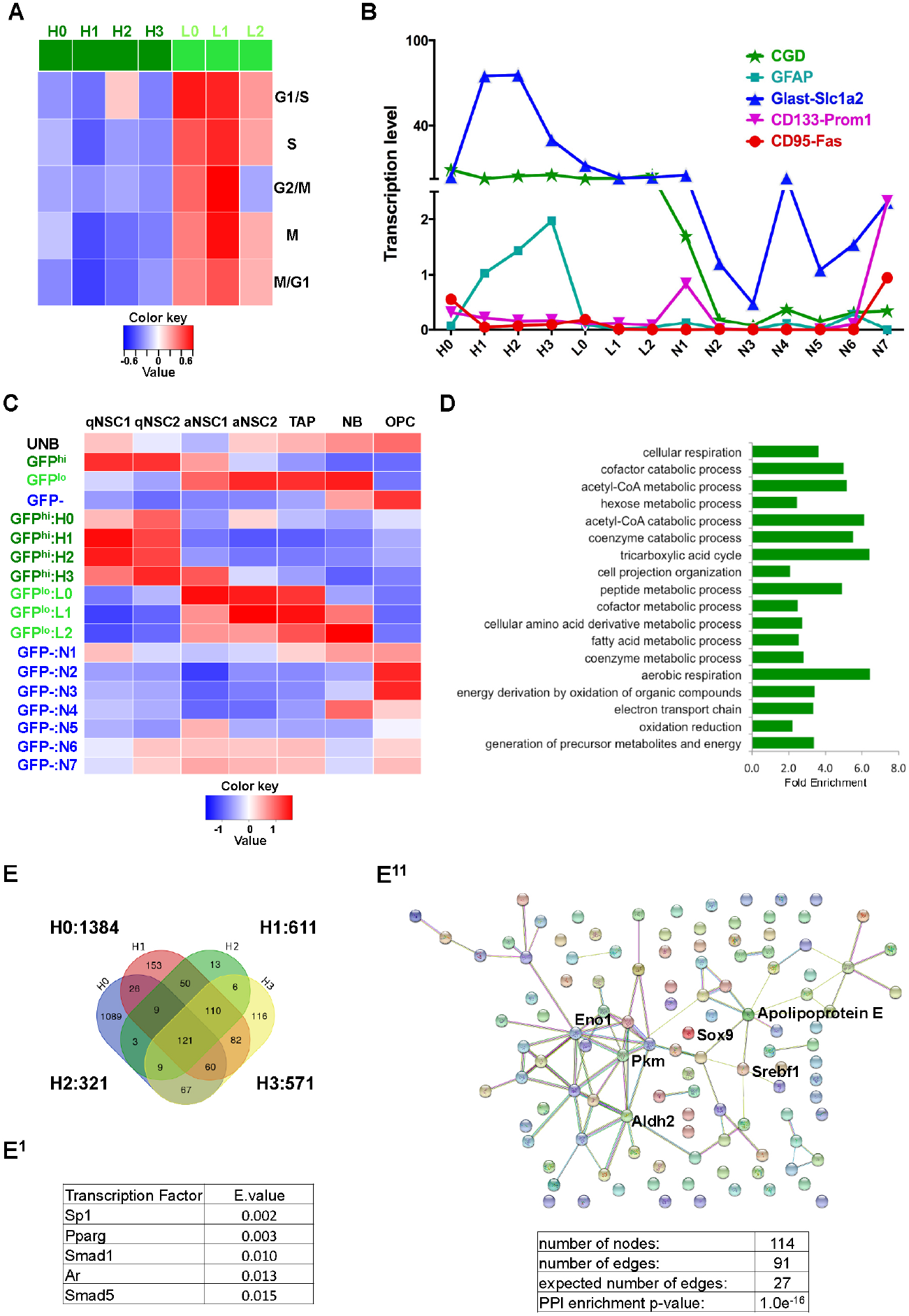
This figure accompanies Figure 4. Granular view provided by analysis of the fourteen groups in the adult SVZ. (A) Cell cycle signature analysis of the GFP^hi^ and GFP^lo^ (H&L) groups reveals heterogeneity. Subgroup L1 shows the highest mitotic index for all seven signatures (see Fig. 2E). (B) Transcription levels of the five NSC markers within the fourteen populations (all standard errors < 1.2, units are normalized values scaled to 10,000 counts/cell). (C) Published single cell gene signatures fail to distinguish between the four GFP^hi^ subgroups. (D) Known and unknown biological processes related to quiescent NSCs are revealed by a comprehensive list of genes derived from H0-H3 groups. (E-E^11^) An overlap analysis of genes commonly expressed in H0-H3 groups results in 121 genes (E), putative transcription factors associated to this list (TfactS analysis) (E^1^), and enrichment for a gene network is revealed (STRING analysis) (E^11^).

**Fig. S5.**
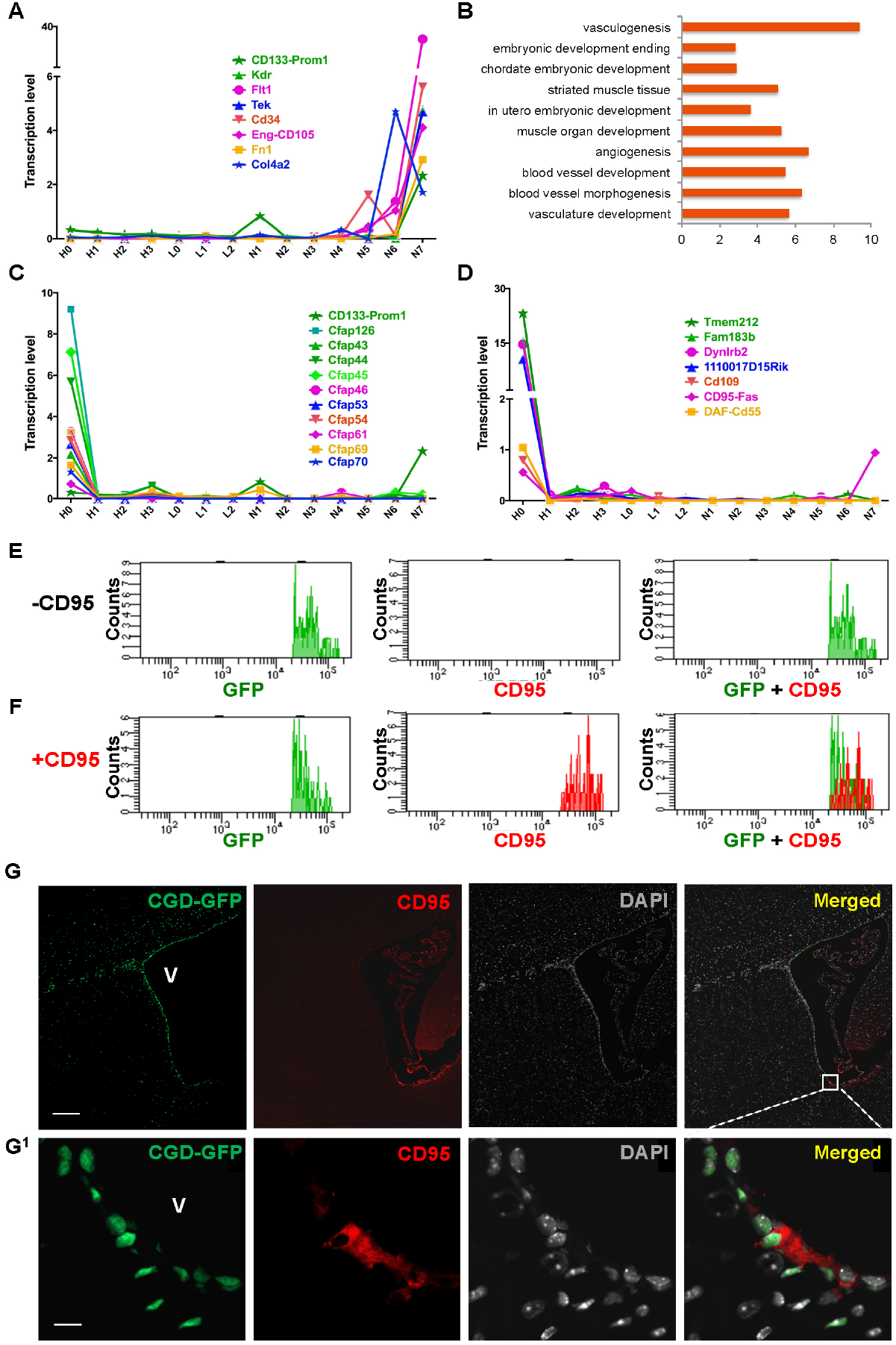
This figure accompanies Figure 5. H0 subgroup gene analysis. (A) A gene network proposed as ependymal-like NSCs is enriched in the GFP-:N7 subgroup that by our analysis represents endothelial progenitor cells (all standard errors < 0.2, units are normalized values scaled to 10,000 counts/cell). (B) Gene ontology analysis of GFP-:N7 associate to vasculature and blood vessel development. (C) Cilia associated genes are uniquely expressed in GFP^hi^:H0 subgroup (all standard errors < 0.2, units are normalized values scaled to 10,000 counts/cell). (D) Expression profiles of seven potential markers for H0 subgroup cells (all standard errors < 2.0, units are normalized values scaled to 10,000 counts/cell). (E) and (F) Among the GFP+ subgroups of cells, CD95 antibody preferentially labels cells with higher levels of GFP proteins. (G) IHC analysis reveals some CD95+ cells co-localized with *CGD*-GFP at ventral SVZ. Scale bars: 200μm (G) and 10μm (G^1^). V: lateral ventricle in (G and G^1^).

**Fig. S6.**
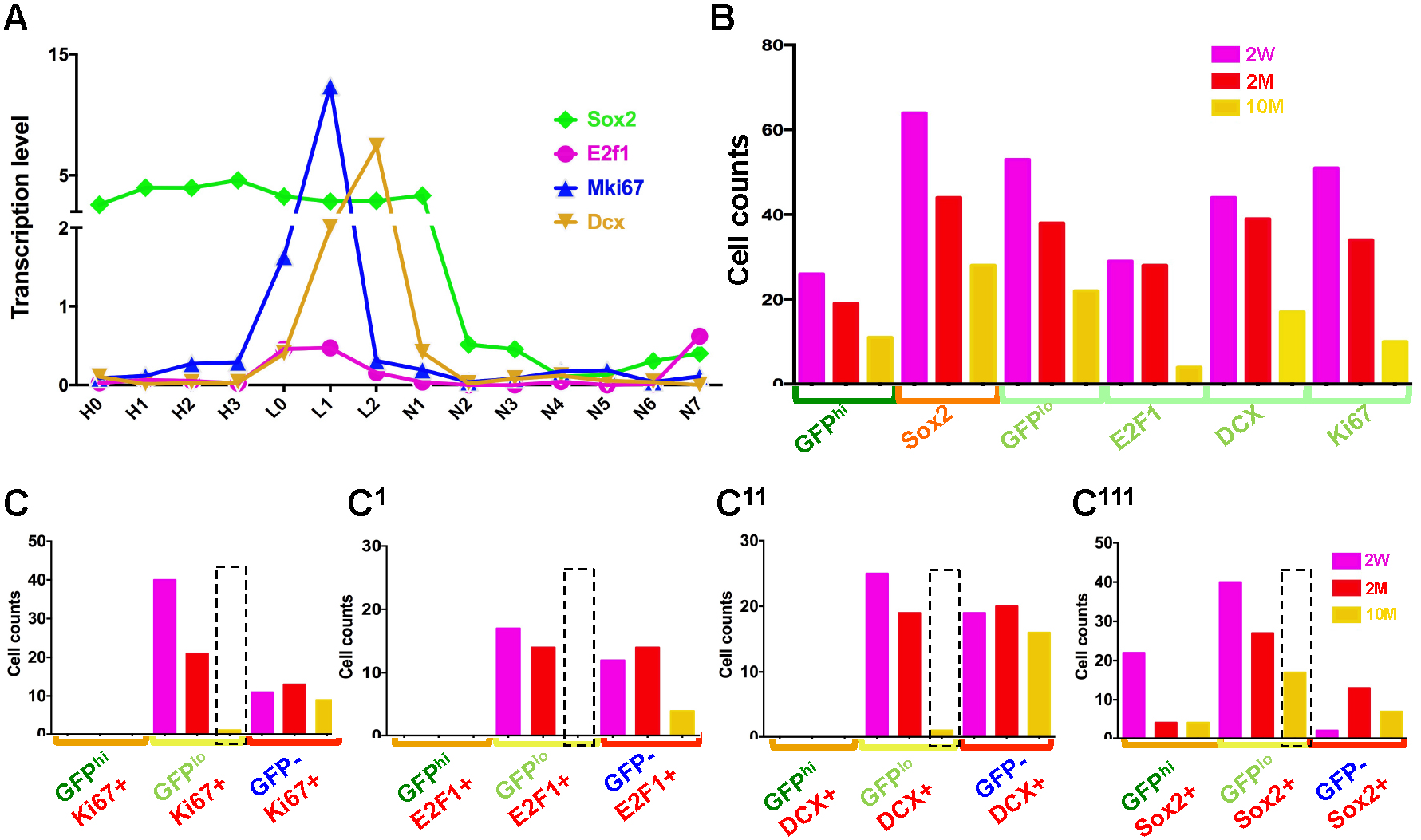
This figure accompanies Figure 6. Age-related alteration of GFP^hi^ and GFP^lo^ lineage related genes. (A) Candidate genes for GFP^hi^ qNSC and GFP^lo^ progenitor cells illustrated in the adult SVZ analysis (all standard errors < 0.6, units are normalized values scaled to 10,000 counts/cell). (B) Concordant with age related GFP expression decrease, cells expressing four candidate genes decrease over age. (C-C^111^) The *CGD-* GFP-associated expression of the three candidate genes for NSC activation (C-C^11^, dashed line) drops more compared to the pan-NSC marker Sox2 (C^111^, dashed line).

**Fig. S7.**
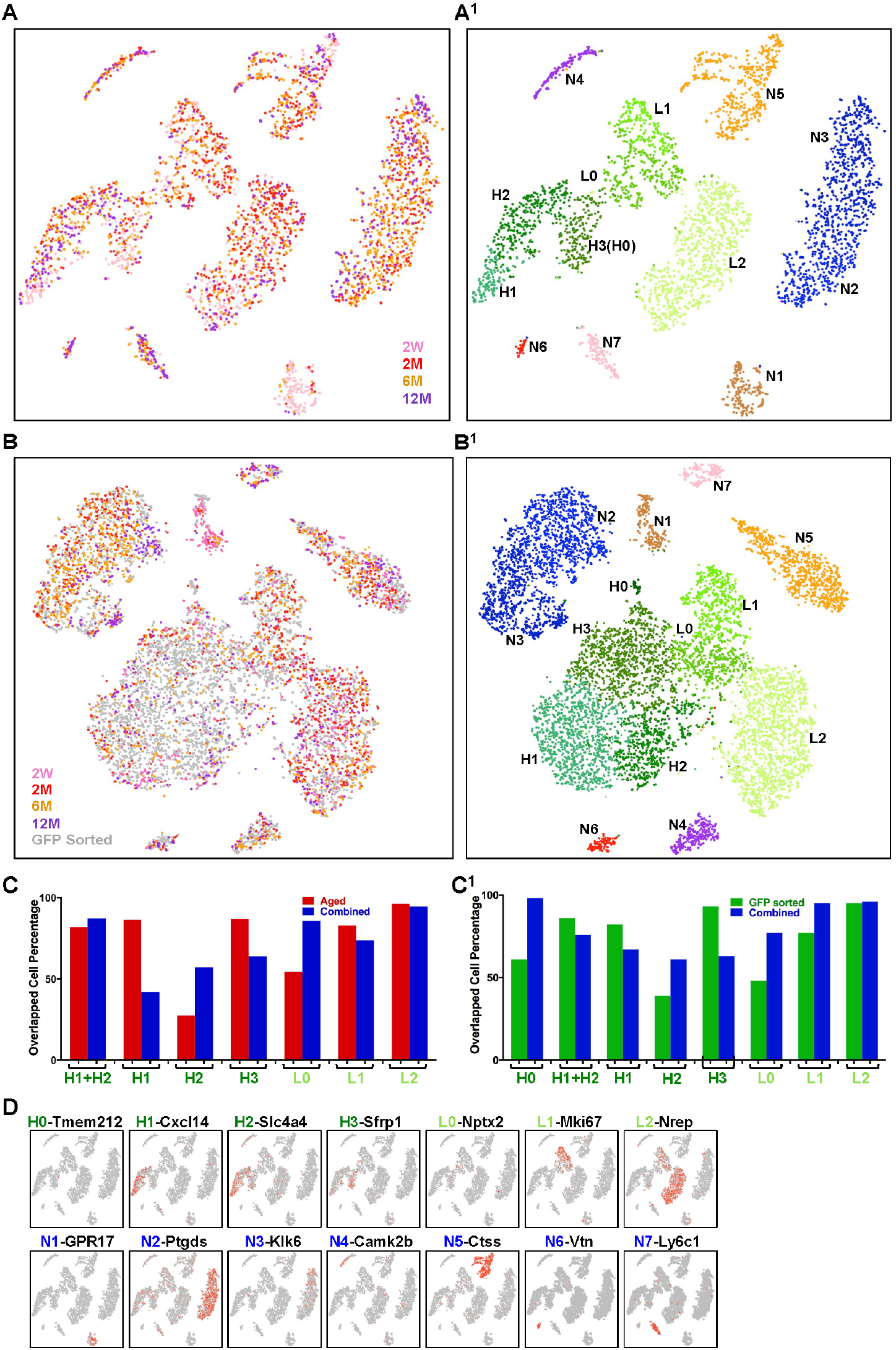
This figure accompanies Figure 7. tSNE projection and marker gene analysis of the SVZ cells over age. (A-A^1^) tSNE projection of the 5,600 single cells from two-week, two-month, six-month, and twelve-month SVZs reveals sample distribution (A) and thirteen groups (A^1^). The groups were assigned by comparing the cell constituents of the GFP sorted cohort (Fig. 3A), the Aged sample cohort (Fig. S7A^1^), and the combined analysis (Fig. S7B^1^). They were further confirmed by comparing unique genes from this analysis (Table S3C) with the unique genes from the GFP sorted cohort analysis (Table S2C). DEGs were used instead when there were not enough unique genes (H2). (B-B^1^) tSNE projection of the comprehensive populations of the combined GFP sorted cohort with the Aged sample cohort (12,200 cells) demonstrates sample distribution (B) and fourteen conserved populations (B^1^). (C-C^1^) Each stem/progenitor group identity in the Aged sample cohort is examined further by comparing the cell constituents in (C) the combined versus the Aged samples and (C^1^) the combined versus the GFP sorted cohort. More than half of the cells are overlapped between the two cohorts in most cases. A significant amount of H2 cells are assigned into the H1 group in the combined analysis, possibly due in part to batch effect correction. (D) Marker genes identified in the two-month SVZ samples guide the identification of Seurat based clusters of SVZ cells from different ages. H0 cells (9 cells in total) are too few to form a group in the Aged cohort by themselves.

## Dataset Table Legends

**Dataset Table S1, related to Figure 2E-2F and S2.** Transcriptomes (TS1A), differentiated expressed genes (TS1B), and GO analysis (TS1C&D) of the bulk RNAseq data derived from the *CGD*-GFP^hi^ vs GFP^lo^ samples. Complete lists of NSC genes derived from various studies including the *CGD*-GFP^hi^ vs GFP^lo^ samples are summarized in TS1E.

**Dataset Table S2, related to Figure 3, S3, 4, S4, and S5A-D**. Transcriptomes (TS2A), differentiated expressed genes (TS2B), unique genes (TS2C), and GO analysis (TS2D) of the fourteen cell groups derived from the two-month old murine SVZs. The 1914 genes derived from H0-H3 groups and associated GOs in Fig. S4D are summarized in (TS2E). The 121 overlapped G genes used in Fig. 4C&S4E, and related transcription factors are shown in (TS2F&G).

**Dataset Table S3, related to Figure 7 and S7**. Transcriptomes (TS3A), differentiated expressed genes (TS3B), and unique genes (TS3C) of the thirteen cell groups derived from the SVZs of two-week, two-month, six-month, and twelve-month mice. Transcriptomes of each cell group in the four aged SVZs are summarized in (TS3D). In lineage H1-H2, genes and GOs up-regulated in either two-month or twelve-month are shown in (TS3E&F). Transcription factors associated to the age-related change are indicated in (TS3G). In lineage H3-L1, genes and GOs up-regulated in either two-month or twelve-month are shown in (TS3H&I). Potential transcription factors involved in the age-related blockage are summarized in (TS3J).

## References

1. Altman J & Das GD (1965) Autoradiographic and histological evidence of postnatal hippocampal neurogenesis in rats. J Comp Neurol 124(3):319–335.

2. Yousefi M, Li L, & Lengner CJ (2017) Hierarchy and Plasticity in the Intestinal Stem Cell Compartment. Trends Cell Biol.

3. Wang LD & Wagers AJ (2011) Dynamic niches in the origination and differentiation of haematopoietic stem cells. Nat Rev Mol Cell Biol 12(10):643–655.

4. Merkle FT, et al. (2014) Adult neural stem cells in distinct microdomains generate previously unknown interneuron types. Nat Neurosci 17(2):207–214.

5. Ming GL & Song H (2011) Adult neurogenesis in the mammalian brain: significant answers and significant questions. Neuron 70(4):687–702.

6. Zeisel A, et al. (2018) Molecular Architecture of the Mouse Nervous System. Cell 174(4):999–1014 e1022.

7. Doetsch F, Caille I, Lim DA, Garcia-Verdugo JM, & Alvarez-Buylla A (1999) Subventricular zone astrocytes are neural stem cells in the adult mammalian brain. Cell 97(6):703–716.

8. Johansson CB, et al. (1999) Identification of a neural stem cell in the adult mammalian central nervous system. Cell 96(1):25–34.

9. Sirko S, et al. (2013) Reactive glia in the injured brain acquire stem cell properties in response to sonic hedgehog. [corrected]. Cell Stem Cell 12(4):426–439.

10. Luo Y, et al. (2015) Single-cell transcriptome analyses reveal signals to activate dormant neural stem cells. Cell 161(5):1175–1186.

11. Lim DA & Alvarez-Buylla A (2014) Adult neural stem cells stake their ground. Trends Neurosci 37(10):563–571.

12. Bjornsson CS, Apostolopoulou M, Tian Y, & Temple S (2015) It takes a village: constructing the neurogenic niche. Dev Cell 32(4):435–446.

13. Jessberger S (2016) Stem Cell-Mediated Regeneration of the Adult Brain. Transfus Med Hemother 43(5):321–326.

14. Dixon KJ, et al. (2015) Endogenous neural stem/progenitor cells stabilize the cortical microenvironment after traumatic brain injury. J Neurotrauma 32(11):753–764.

15. Alcantara Llaguno S, et al. (2009) Malignant astrocytomas originate from neural stem/progenitor cells in a somatic tumor suppressor mouse model. Cancer Cell 15(1):45–56.

16. Li P, et al. (2013) A population of Nestin-expressing progenitors in the cerebellum exhibits increased tumorigenicity. Nat Neurosci 16(12):1737–1744.

17. Alcantara Llaguno SR, et al. (2015) Adult Lineage-Restricted CNS Progenitors Specify Distinct Glioblastoma Subtypes. Cancer Cell 28(4):429–440.

18. Alcantara Llaguno S, et al. (2019) Cell-of-origin susceptibility to glioblastoma formation declines with neural lineage restriction. Nat Neurosci 22(4):545–555.

19. Coskun V, et al. (2008) CD133+ neural stem cells in the ependyma of mammalian postnatal forebrain. Proc Natl Acad Sci U S A 105(3):1026–1031.

20. Beckervordersandforth R, et al. (2010) In vivo fate mapping and expression analysis reveals molecular hallmarks of prospectively isolated adult neural stem cells. Cell Stem Cell 7(6):744–758.

21. Codega P, et al. (2014) Prospective identification and purification of quiescent adult neural stem cells from their in vivo niche. Neuron 82(3):545–559.

22. Mich JK, et al. (2014) Prospective identification of functionally distinct stem cells and neurosphere-initiating cells in adult mouse forebrain. Elife 3:e02669.

23. Llorens-Bobadilla E, et al. (2015) Single-Cell Transcriptomics Reveals a Population of Dormant Neural Stem Cells that Become Activated upon Brain Injury. Cell Stem Cell 17(3):329–340.

24. Dulken BW, Leeman DS, Boutet SC, Hebestreit K, & Brunet A (2017) Single-Cell Transcriptomic Analysis Defines Heterogeneity and Transcriptional Dynamics in the Adult Neural Stem Cell Lineage. Cell Rep 18(3):777–790.

25. Zimmerman L, et al. (1994) Independent regulatory elements in the nestin gene direct transgene expression to neural stem cells or muscle precursors. Neuron 12(1):11–24.

26. Mignone JL, Kukekov V, Chiang AS, Steindler D, & Enikolopov G (2004) Neural stem and progenitor cells in nestin-GFP transgenic mice. J Comp Neurol 469(3):311–324.

27. Imayoshi I, Ohtsuka T, Metzger D, Chambon P, & Kageyama R (2006) Temporal regulation of Cre recombinase activity in neural stem cells. Genesis 44(5):233–238.

28. Yu TS, Zhang G, Liebl DJ, & Kernie SG (2008) Traumatic brain injury-induced hippocampal neurogenesis requires activation of early nestinexpressing progenitors. J Neurosci 28(48):12901–12912.

29. Chen J, Kwon CH, Lin L, Li Y, & Parada LF (2008) Inducible site-specific recombination in neural stem/progenitor cells. submitted.

30. Zywitza V, Misios A, Bunatyan L, Willnow TE, & Rajewsky N (2018) Single-Cell Transcriptomics Characterizes Cell Types in the Subventricular Zone and Uncovers Molecular Defects Impairing Adult Neurogenesis. Cell Rep 25(9):2457–2469 e2458.

31. Mizrak D, et al. (2019) Single-Cell Analysis of Regional Differences in Adult V-SVZ Neural Stem Cell Lineages. Cell Rep 26(2):394–406 e395.

32. Ninkovic J, Mori T, & Gotz M (2007) Distinct modes of neuron addition in adult mouse neurogenesis. J Neurosci 27(40):10906–10911.

33. Mirzadeh Z, Merkle FT, Soriano-Navarro M, Garcia-Verdugo JM, & Alvarez-Buylla A (2008) Neural stem cells confer unique pinwheel architecture to the ventricular surface in neurogenic regions of the adult brain. Cell Stem Cell 3(3):265–278.

34. Doetsch F, Garcia-Verdugo JM, & Alvarez-Buylla A (1997) Cellular composition and three-dimensional organization of the subventricular germinal zone in the adult mammalian brain. J Neurosci 17(13):5046–5061.

35. Whitfield ML, et al. (2002) Identification of genes periodically expressed in the human cell cycle and their expression in tumors. Mol Biol Cell 13(6):1977–2000.

36. Subramanian A, Kuehn H, Gould J, Tamayo P, & Mesirov JP (2007) GSEA-P: a desktop application for Gene Set Enrichment Analysis. Bioinformatics 23(23):3251–3253.

37. Macosko EZ, et al. (2015) Highly Parallel Genome-wide Expression Profiling of Individual Cells Using Nanoliter Droplets. Cell 161(5):1202–1214.

38. Huang da W, Sherman BT, & Lempicki RA (2009) Systematic and integrative analysis of large gene lists using DAVID bioinformatics resources. Nat Protoc 4(1):44–57.

39. Angerer P, et al. (2016) destiny: diffusion maps for large-scale single-cell data in R. Bioinformatics 32(8):1241–1243.

40. Fiorelli R, Azim K, Fischer B, & Raineteau O (2015) Adding a spatial dimension to postnatal ventricular-subventricular zone neurogenesis. Development 142(12):2109–2120.

41. Lao CL, Lu CS, & Chen JC (2013) Dopamine D3 receptor activation promotes neural stem/progenitor cell proliferation through AKT and ERK1/2 pathways and expands type-B and -C cells in adult subventricular zone. Glia 61(4):475–489.

42. Shah PT, et al. (2018) Single-Cell Transcriptomics and Fate Mapping of Ependymal Cells Reveals an Absence of Neural Stem Cell Function. Cell 173(4):1045–1057 e1049.

43. Roman-Trufero M, et al. (2009) Maintenance of undifferentiated state and self-renewal of embryonic neural stem cells by Polycomb protein Ring1B. Stem Cells 27(7): 1559–1570.

44. Delgado AC, et al. (2014) Endothelial NT-3 delivered by vasculature and CSF promotes quiescence of subependymal neural stem cells through nitric oxide induction. Neuron 83(3):572–585.

45. Reynolds BA & Rietze RL (2005) Neural stem cells and neurospheres--re-evaluating the relationship. Nat Methods 2(5):333–336.

46. Pastrana E, Silva-Vargas V, & Doetsch F (2011) Eyes wide open: a critical review of sphere-formation as an assay for stem cells. Cell Stem Cell 8(5):486–498.

47. Belenguer G, Domingo-Muelas A, Ferron SR, Morante-Redolat JM, & Farinas I (2016) Isolation, culture and analysis of adult subependymal neural stem cells. Differentiation 91(4-5):28–41.

48. Enwere E, et al. (2004) Aging results in reduced epidermal growth factor receptor signaling, diminished olfactory neurogenesis, and deficits in fine olfactory discrimination. J Neurosci 24(38):8354–8365.

49. Shi Z, et al. (2017) Single-cell transcriptomics reveals gene signatures and alterations associated with aging in distinct neural stem/progenitor cell subpopulations. Protein Cell.

50. Leeman DS, et al. (2018) Lysosome activation clears aggregates and enhances quiescent neural stem cell activation during aging. Science 359(6381):1277–1283.

51. Daynac M, Morizur L, Chicheportiche A, Mouthon MA, & Boussin FD (2016) Age-related neurogenesis decline in the subventricular zone is associated with specific cell cycle regulation changes in activated neural stem cells. Sci Rep 6:21505.

52. Mizuguchi H, Xu Z, Ishii-Watabe A, Uchida E, & Hayakawa T (2000) IRES-dependent second gene expression is significantly lower than capdependent first gene expression in a bicistronic vector. Mol Ther 1(4):376–382.

53. Szymczak AL, et al. (2004) Correction of multi-gene deficiency in vivo using a single ‘self-cleaving’ 2A peptide-based retroviral vector. Nat Biotechnol 22(5):589–594.

54. Kim JH, et al. (2011) High cleavage efficiency of a 2A peptide derived from porcine teschovirus-1 in human cell lines, zebrafish and mice. PLoS One 6(4):e18556.

55. Mirzadeh Z, Doetsch F, Sawamoto K, Wichterle H, & Alvarez-Buylla A (2010) The subventricular zone en-face: wholemount staining and ependymal flow. J Vis Exp (39).

56. Ferron SR, et al. (2007) A combined ex/in vivo assay to detect effects of exogenously added factors in neural stem cells. Nat Protoc 2(4):849–859.

## SI References

1. Macosko EZ, et al. (2015) Highly Parallel Genome-wide Expression Profiling of Individual Cells Using Nanoliter Droplets. Cell 161(5):1202–1214.

2. Huang da W, Sherman BT, & Lempicki RA (2009) Systematic and integrative analysis of large gene lists using DAVID bioinformatics resources. Nat Protoc 4(1):44–57.

3. Angerer P, et al. (2016) destiny: diffusion maps for large-scale single-cell data in R. Bioinformatics 32(8):1241–1243.

4. Lein ES, et al. (2007) Genome-wide atlas of gene expression in the adult mouse brain. Nature 445(7124):168–176.

5. Heberle H, Meirelles GV, da Silva FR, Telles GP, & Minghim R (2015) InteractiVenn: a web-based tool for the analysis of sets through Venn diagrams. BMC Bioinformatics 16:169.

6. Essaghir A, et al. (2010) Transcription factor regulation can be accurately predicted from the presence of target gene signatures in microarray gene expression data. Nucleic Acids Res 38(11):e120.

7. Szklarczyk D, et al. (2017) The STRING database in 2017: quality-controlled protein-protein association networks, made broadly accessible. Nucleic Acids Res 45(D1):D362–D368.

8. Wang X, Spandidos A, Wang H, & Seed B (2012) PrimerBank: a PCR primer database for quantitative gene expression analysis, 2012 update. Nucleic Acids Res 40(Database issue):D1144–1149.

9. Sabbagha NG, et al. (2011) Alternative splicing in Acad8 resulting a mitochondrial defect and progressive hepatic steatosis in mice. Pediatr Res 70(1):31–36.

10. Kharchenko PV, Silberstein L, & Scadden DT (2014) Bayesian approach to single-cell differential expression analysis. Nat Methods 11(7):740–742.

